# Molecular basis of C9orf72 poly-PR interference with the β-karyopherin family of nuclear transport receptors

**DOI:** 10.1101/2022.05.05.490606

**Authors:** Hamidreza Jafarinia, Erik Van der Giessen, Patrick R. Onck

## Abstract

Nucleocytoplasmic transport (NCT) is affected in several neurodegenerative diseases including C9orf72-ALS. It has recently been found that arginine-containing dipeptide repeat proteins (R-DPRs), translated from C9orf72 repeat expansions, directly bind to several importins. To gain insight into how this can affect nucleocytoplasmic transport, we use coarse-grained molecular dynamics simulations to study the molecular interaction of poly-PR, the most toxic DPR, with several Kapβs (importins and exportins). We show that poly-PR–Kapβ binding depends on the net charge per residue (NCPR) of the Kapβ, salt concentration of the solvent, and poly-PR length. Poly-PR makes contact with the inner surface of most importins, which strongly interferes with Kapβ binding to cargo-NLS, IBB, and RanGTP in a poly-PR length-dependent manner. Longer poly-PRs at higher concentrations are also able to make contact with the outer surface of importins that contain several binding sites to FG-Nups. We also show that poly-PR binds to exportins, especially at lower salt concentrations, interacting with several RanGTP and FG-Nup binding sites. Overall, our results suggest that poly-PR might cause length-dependent defects in cargo loading, cargo release, Kapβ transport and Ran gradient across the nuclear envelope.

## Introduction

The G4C2 hexanucleotide repeat expansion in C9orf72 is the most frequent genetic cause of amyotrophic lateral sclerosis (ALS) and frontotemporal dementia (FTD) [1, 2]. Healthy individuals typically have up to around 20 repeats of G4C2, while patients with C9orf72-mediated ALS/FTD (C9-ALS/FTD) usually have hundreds to thousands repeats [1-3]. The RNA transcripts of this repeat expansion are translated to five types of dipeptide repeat proteins (DPRs): poly-PR, poly-GR, poly-GA, poly-GP, and poly-PA [4, 5]. Toxic gain of function of RNA transcripts [6, 7] and DPRs [4, 5, 8], and loss of function of the C9orf72 protein [1, 9] are thought to cause C9-ALS/FTD. Among the DPRs, the positively-charged arginine-containing DPRs (R-DPRs) show the highest levels of toxicity in both cell and animal models [8, 10-16].

R-DPRs are hypothesized to cause a wide variety of cellular defects [17]. Recent evidence suggests a link between R-DPRs and defects in the transport of macromolecular cargoes between nucleus and cytoplasm [18-20]. This nucleocytoplasmic transport (NCT) [21, 22] is mediated by large multiprotein assemblies called nuclear pore complexes (NPCs). The central channel of the NPC is lined with intrinsically disordered phenylalanine-glycine-rich Nups (FG-Nups) that collectively function as a selective permeability barrier of the NPC. This FG-barrier allows the passive diffusion of small molecules below ∼30-40 kDa, while it increasingly slows down the passage of larger cargoes unless they are bound to nuclear transport receptors (NTRs) [23-25]. The β-karyopherin (Kapβ) family is the largest class of NTRs that includes both import and export receptors [26]. Importins transport nuclear localization signal (NLS)-containing cargoes into the nucleus, and exportins transport nuclear export signal (NES)-containing cargoes out of the nucleus. NTRs can either bind directly to an NLS/NES-containing cargo (referred to as NLS/NES-cargo), or, in case of importin β1 (Impβ1), bind indirectly through the adaptor proteins importin α (Impα) or snurportin-1 (SPN1) [27-29]. The adaptor proteins bind to Impβ1 through their N-terminal importin β-binding (IBB) domains. Another essential ingredient in NCT is the GTPase Ran which switches between guanosine triphosphate (GTP)- and guanosine diphosphate (GDP)-bound forms [30]. The directionality of NCT is mediated via a steep RanGTP-RanGDP gradient over the nuclear envelope [31]. In the import cycle, RanGTP disassembles the importin-NLS-cargo complex in the nucleus by binding to the importin, and the resulting RanGTP-importin complex shuttles back to the cytoplasm. In the export process, RanGTP promotes the formation of a RanGTP-exportin-NES-cargo complex in the nucleus which facilitates the export of the NES-cargo to the cytoplasm. The hydrolysis of RanGTP to RanGDP in the cytoplasm disassembles the RanGTP-importin complex in the import cycle and the RanGTP-exportin-NES-cargo complex in the export cycle [30].

Several studies have reported NTRs as potential interactors of R-DPRs [13, 20, 32]. Recently, R-DPRs with 25 repeat units have been shown to directly bind to several importins: Impβ1, Imp5, Imp7, Imp9, TNPO1, and TNPO3, but not to exportin CRM1 in *in vitro* experiments [19]. R-DPRs (with 10 or 25 repeat units) have also been found to cause defects in Impβ1- and TNPO1-mediated import [19, 20]. Although these studies provide interesting insights into potential pathways for R-DPRs-mediated NCT defects, the molecular basis for the interaction of longer R-DPRs with different Kapβs and the resulting NCT defects has not yet been identified. In this study, we use coarse-grained (CG) molecular dynamics models [33-37] to investigate the length-dependent interaction of poly-PR (known to be the most toxic DPR [11-13]) with different members of the Kapβ family of NTRs with the aim to provide mechanistic insight into the way poly-PR interferes with the transport functionality of Kapβs.

## Results and discussion

### Coarse-grained models of poly-PR and Kapβs

We use a CG modeling framework to investigate the interaction between poly-PR and several members of the Kapβ family of NTRs in their free, unbound state. In this study, we consider human Kapβs: importins Impβ1, Imp5, TNPO1, TNPO3, and exportins Exp1 (also known as CRM1), Exp5 (also known as XPO5), as well as yeast Kapβs: importins KAP95, KAP114, KAP120, KAP121, and exportin KAP124. The selected yeast Kapβs are homologs of human Impβ1, Imp9, Imp11, Imp5, and CRM1, respectively [30, 38].

Our simulations adopt one-bead-per-amino-acid (1BPA) CG models of poly-PR and the selected Kapβs, see figure 1a,b. In this approach, initially designed to represent the FG-nups in the NPC [33, 39], each residue is represented by a single bead located at the position of the α-carbon atom. The CG model of poly-PR, see figure 1a (left panel), has already been applied in a previous study on the length-dependent phase separation of poly-PR and negatively-charged peptides [35]. Kapβs consist of tandem HEAT repeats, arranged into superhelical structures in case of importins and ring-shaped/U-shaped structures in case of exportins [40-42]. Each HEAT repeat contains two antiparallel alpha helices, referred to as A- and B-helices, connected by loops of varying length [42]. In figure 1a (right panel), some α-carbon beads are shown on top of the crystal structure of Impβ1. The crystal structures used to build residue-scale CG models of Kapβs, see figure 1b, are summarized in table S3 in the SI. The overall structure of Kapβs is preserved using a network of stiff harmonic bonds. In addition, the distribution of charged and aromatic residues that are relevant for the interaction of R-DPRs with NTRs are included in the model. Our implicit-solvent CG force field also accounts for the screening effect of ions. More details about the CG modeling and force field are provided in the Methods section and section 1 of the SI.

**Figure 1:**
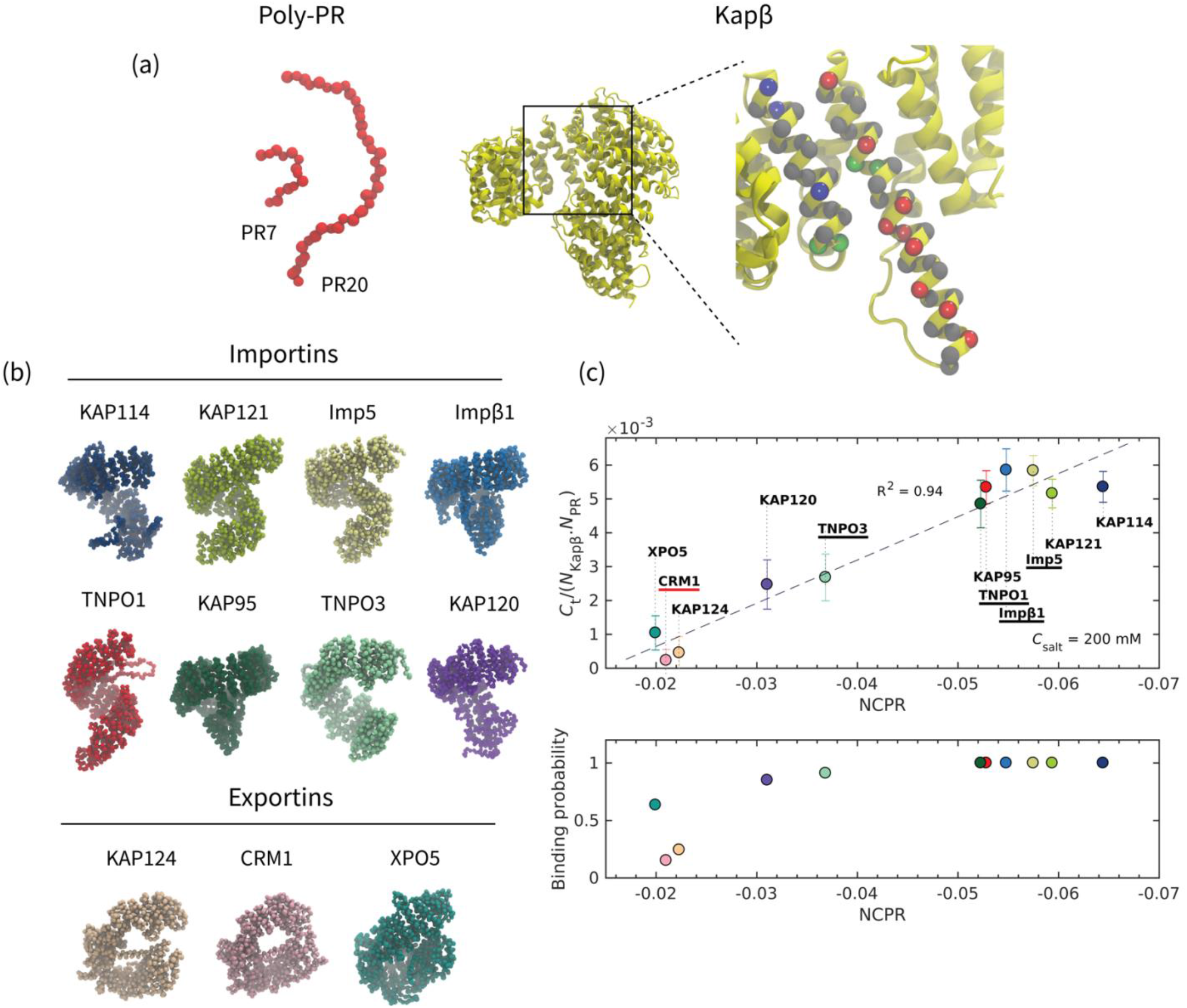
Coarse-grained modeling shows the importance of electrostatic interactions for the binding between poly-PR and Kapβs. (a) (Left panel) 1-bead-per-amino-acid (1BPA) representation of poly-PR with 7 and 20 repeat units. Both Proline and Arginine residues are shown in red. (Right panel) Our 1BPA approach to create coarse-grained (CG) models for Kapβs. The CG beads are placed at the location of α-carbon atoms, here shown for two α-helices of impβ1 on top of the crystal structure. Red, blue, and green beads correspond to negatively-charged residues (D and E), positively-charged residues (R and K), and residues with aromatic rings (F, Y, and W), respectively. Other residues are shown with grey beads. The all-atom crystal structure is depicted in yellow using the New Cartoon representation in VMD. (b) 1BPA representation of the various Kapβs modeled in the current study that serve either as importin or exportin. Importins have superhelical structures whereas exportins have a more ring-like structure in case of KAP124 and CRM1, and U-shaped structure in case of XPO5. The crystal structures used to make the CG models are listed in table S3. (c) (Top panel) The time-averaged number of contacts *C*_*t*_ between PR25 and Kapβs plotted against the net charge per residue (NCPR) of Kapβs at a monovalent salt concentration *C*_salt_ of 200 mM. The values on the vertical axis are normalized by the sequence length of the importins/exportins (*N*_Kapβ_) and the sequence length of poly-PR (*N*_PR_). The NCPR values on the horizontal axis is calculated based on the sequence lengths and amino acid compositions of the Kapβ models listed in table S3 (column 6) and table S5. Among the selected Kapβs, the importins Imp5, Impβ1, TNPO1, TNPO3 (highlighted with a horizontal black line) have been shown to directly bind to PR25 *in vitro* experiments [19]. However, no binding has been observed for CRM1 (highlighted with a horizontal red line) in *in vitro* experiments performed at the same monovalent salt concentration [19]. A linear correlation can be seen between the normalized *C*_*t*_ and the negative NCPR. The dashed line shows a linear fit. The error bars are half of the standard deviation (see the SI for details). (Bottom panel) The PR25-Kapβs binding probability P_b_ plotted against NCPR of Kapβs at *C*_salt_ = 200 mM.

### Poly-PR interaction with Kapβs correlates with the net charge per residue (NCPR) of Kapβs

Among the Kapβs shown in figure 1b, TNPO1, TNPO3, Imp5, and Imp*β* have been shown to directly bind to PR25 (poly-PR with 25 PR repeat units), and no binding has been observed for CRM1 *in vitro* [19]. In figure 1c, we show our simulation results for PR25 interacting with the selected Kapβs. In order to allow for a comparison with experimental findings, simulations in this section are performed at a monovalent salt concentration *C*_salt_ = 200 mM. In each simulation, the periodic simulation box contains one copy of PR25 and one copy of a Kapβ. To quantify the interaction between PR25 and Kapβs, we calculate (i) the time-averaged number of contacts *C*_*t*_ between PR25 and each Kapβ using a cutoff of 1 nm, and (ii) the binding probability, taken as the probability of having more than 10% of the poly-PR residues within 1 nm proximity of each Kapβ. In figure 1c, the values of *C*_*t*_ and the binding probability are plotted against the net charge per residue (NCPR) of the Kapβs, with NCPR being the total charge of the protein (in units of elementary charge) divided by its sequence length (see tables S3 and S5 for the sequence lengths and amino acid compositions of the Kapβ models used). The values of *C*_*t*_ are normalized by the sequence length of the importins/exportins (*N*_Kapβ_) and the sequence length of poly-PR (*N*_PR_). Our results in figure 1c (top panel), show a linear correlation between the normalized number of contacts *C*_*t*_ and the negative NCPR of the Kapβs. PR25 makes more contacts with the importins, especially KAP95, TNPO1, Impβ1, Imp5, KAP121, and KAP114, than with the three exportins, XPO5, CRM1, and KAP124, due to the difference in negative NCPR. No such correlation is observed between *C*_*t*_ and the aromatic residue content (data not shown). The results also reveal high binding probabilities (> 0.85) for importins, and lower values for the exportins. CRM1 shows the lowest binding probability (∼0.15), consistent with the absence of binding in experiments [19]. We show in figure S2 of the SI that varying the binding criterion does not affect the binding probabilities for Kapβs with high binding probabilities (∼1), and only slightly changes the binding probabilities for other Kapβs. Overall, our results show that the poly-PR interaction with Kapβs is mostly driven by electrostatic interactions, in correspondence with *in vitro* binding assays.

### Salt concentration and poly-PR length affect the poly-PR–Kapβ binding behavior

It has been found that the toxicity of poly-PR increases with the number of PR repeat units [8, 11, 43-45], but this has so far not been linked to the interaction of poly-PR with Kapβs. To provide insight into this, we study the interaction of one copy of poly-PR that has either 7, 20, 35, or 50 repeat units with one copy of each Kapβ. To quantify the interaction, the normalized number of contacts *C*_*t*_/*(N*_Kapβ_*N*_PR_) and binding probability are reported in figure 2a. The simulations are performed at two monovalent salt concentrations: *C*_salt_ = 200 mM, as in the previous section and similar to previous *in vitro* experiments, and *C*_salt_ = 100 mM, corresponding to that of the cytoplasm [46]. Increasing the poly-PR length increases the number of contacts *C*_*t*_ (only the normalized *C*_*t*_ is shown), and the binding probability (figure 2a, bottom panel). The effect of the poly-PR length on the binding probability is much more pronounced at *C*_salt_ = 200 mM. The reason for this is that reduction of the salt concentration to 100 mM reduces screening effects and thus increases the binding probability, thus confirming the important role of electrostatic interactions in poly-PR–Kapβ interactions. This increased binding probability is especially visible for the interaction of shorter poly-PRs with Kapβs that have lower negative NCPR. At *C*_salt_ = 100 mM, poly-PR with more than 20 repeat units also bind to CRM1 and Kap124 with binding probabilities ∼1, showing that a reduction of the salt concentration can significantly increase the binding probability for these two exportins. As can be seen in figure 2a, when the poly-PR–Kapβ binding probability is high (∼1) for a certain Kapβ, i.e. the same *N*_Kapβ_, a shorter poly-PR makes more contacts per unit length (*C*_*t*_/*N*_PR_) with the Kapβ compared to a longer poly-PR. When a relatively short poly-PR chain (for example PR7 or PR20) binds to a Kapβ, almost all its residues make contact with the Kapβ. However, in the bound states of a relatively long poly-PR (for example PR50), some regions make less or no contact with the Kapβ thus resulting in a lower time-averaged number of contacts per unit length of poly-PR. The snapshots of figure 2b clearly illustrate this difference in binding of PR7 and PR50 to Impβ1.

**Figure 2:**
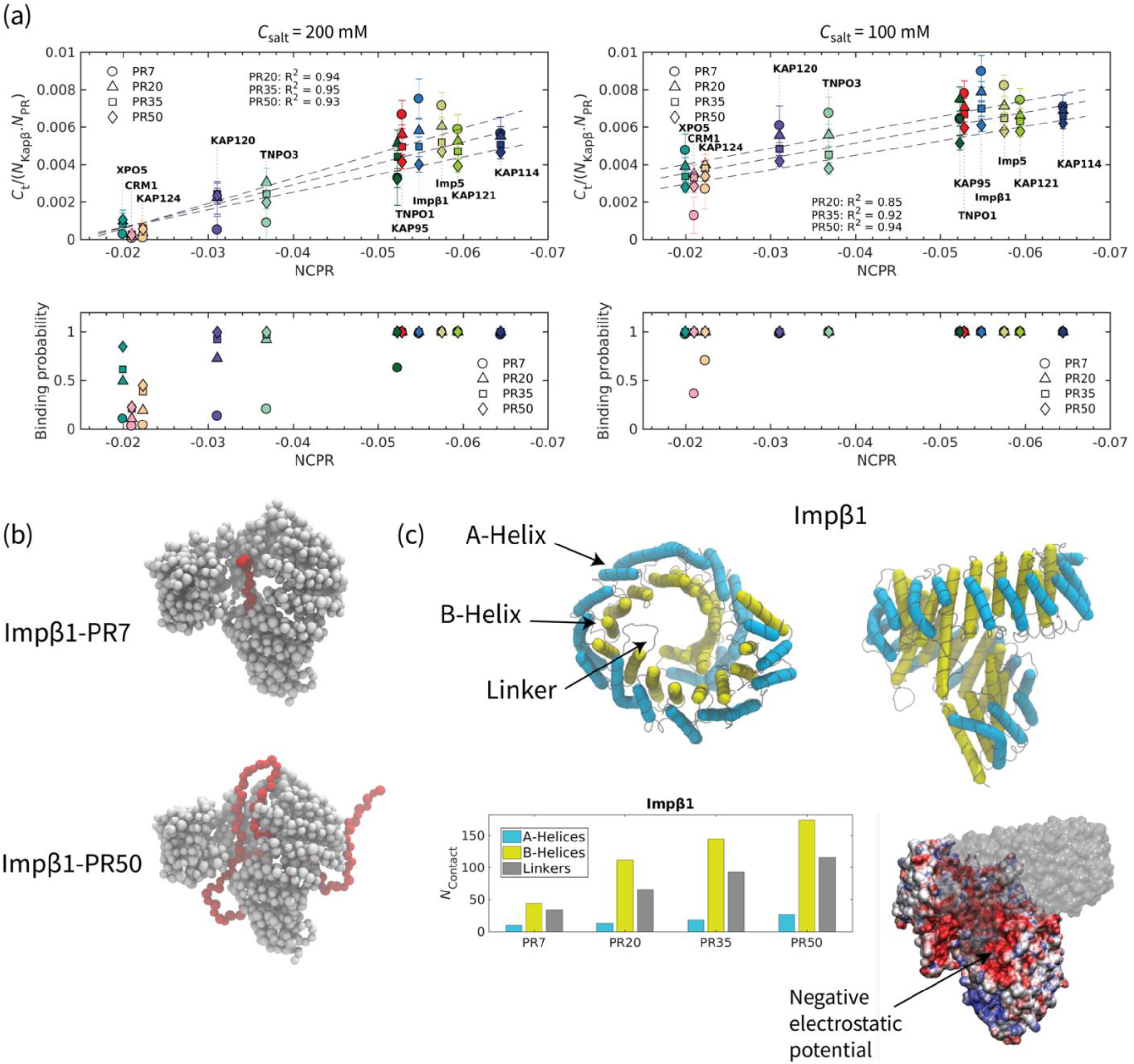
Salt concentration and poly-PR length affect the poly-PR–Kapβ binding behavior. (a) Normalized time-average number of contacts *C*_*t*_, and binding probability P_b_ for the interaction between poly-PR with 7, 20, 25 and 50 repeat units and Kapβs at monovalent salt concentrations of *C*_salt_ = 200 mM (left panel) and *C*_salt_ = 100 mM (right panel). A linear correlation can be seen between the normalized *C*_*t*_ and the negative NCPR of Kapβs for poly-PR with number of repeat units ≥ 20. The dashed lines show linear fits. The error bars are half of the standard deviation. The data for PR50 interaction with KAP120 and KAP114 were taken from [50]. (b) Sample snapshots showing the binding of PR7 and PR50 to Impβ1. In each snapshot poly-PR is depicted in red and Impβ1 is depicted in light grey. (c) (Top panels) The structure of Impβ1 shown from a top view (left) along the superhelical axis and a side view (right). A- and B-helices of Impβ1 are highlighted with light blue and yellow tubes, respectively, using the Bendix plugin in VMD. The linkers that connect the A- and B-helices are shown in light grey. A-helices constitute the inner surface and B-helices constitute the other surface of Impβ1. See table S4 for snapshots of the other Kapβs used in this study. (Bottom panels) The number of residues in each region of Impβ that make contact with poly-PR (*N*_contact_) at *C*_salt_ = 100 plotted for PR7, PR20, PR35 and PR50 (for details see section 2 of the SI). The results for other Kapβs are shown in figure S4. Poly-PR tend to make more contacts with the inner surface of Impβ1, i.e. B-helices. The electrostatic potential of Impβ1 shows negatively-charged regions (shown with an arrow) on the inner surface of Impβ1. The electrostatic surface potentials are obtained using PDB2PQR [51] and plotted using the Surf representation in VMD on a red-white-blue map. Positive and negative surface potentials are denoted by blue and red, respectively. For better visualization, part of the electrostatic potential surface is depicted using a transparent surface. Electrostatic potentials for the other Kapβs are presented in table S4.

To study in more detail how poly-PR targets Kapβs, we first identify the A-helices, B-helices and linkers in the crystal structures of Kapβs (see figure 2c), using the STRIDE secondary structure prediction algorithm [47] (for details, see section 3 of the SI). The linker regions contain residues that connect A- and B-helices in each HEAT repeat as well as the residues that connect consecutive HEAT repeats. In figure 2c, the A- and B-helices for all 19 HEAT repeats of Impβ1 are highlighted. For the other Kapβs, see the snapshots in table S4 in the SI. In general, the A-helices form the outer convex surface and the B-helices form the inner concave surface of Impβ1. In the bottom panel of figure 2c, we report *N*_contact_, the number of residues in the A-helices, B-helices, and linkers of Impβ1 that make contact with poly-PR at *C*_salt_ of 100 mM. For the calculation of *N*_contact_, a Kapβ residue is considered to be a contact site if the contact probability for this residue is larger than 0.10 (for details see section 2 of the SI). Longer poly-PRs are seen to make more contact with Impβ1 residues. The results in figure 2c also show that poly-PR interacts more with the B-helices and the linkers than with the A-helices. The larger number of contacts with the B-helices can be explained by the higher negative electrostatic potential of the inner surface of Impβ1, see figure 2c bottom right. The same binding behavior can also be seen for other importins: KAP114, KAP121, Imp5, TNPO1, KAP95, and TNPO3, see the left column of figure S4 in the SI. The result for TNPO1 is consistent with the binding of poly-PR to a negatively-charged cavity in the inner surface of the TNPO1 [48]. Snapshots presented in table S4 of the SI show that all the importins contain regions with a relatively high negative electrostatic potential at their inner surface. The results in figure S4 and S5 show that longer poly-PRs are also able to interact with the outer surface of the importins because increasing poly-PR length increases the number of contact residues in the A-helices. This effect is most pronounced for Imp5, Impβ1, TNPO1, KAP95, and TNPO3.

For the exportins, on the other hand, we observe different binding behavior with a more dominant role for the linker regions. For poly-PR with length ≤ 20 repeat units interacting with CRM1 and its yeast homolog KAP124, we observe a slightly higher *N*_contact_ for the B-helices. However, longer poly-PRs interact equally with both A- and B-helices of these two exportins, see right column of figure S4. This can be due to a less significant difference between the electrostatic potential of the inner and outer surfaces of these exportins compared to importins. The interaction of XPO5 is different from the other exportins and importins as in this case the poly-PR mostly interact with the linkers that connect the A and B-helices. XPO5 has a closed U-shaped conformation in its cargo-free state, and in contrast to other Kapβs, the inner surface of XPO5 is positively charged and used for the binding and export of miRNA [49]. A more detailed analysis of the contact sites will be provided in the following section.

### Poly-PR interacts with NLS-cargo, IBB, RanGTP and FG-Nup binding sites of different Kapβs in a poly-PR length-dependent manner

To provide more molecular insight into the way poly-PR target each Kapβ, we investigate the contact probability of each Kapβ residue in interacting with poly-PR (for more details about contact probability see the Methods section and section 2 of the SI). In figure 3a, we show the results for the interaction of PR7 and PR50 with the importin Impβ1 and the exportin Kap124. Results for the other Kapβs can be found in figure S6. For comparison, we also highlight the known NLS-cargo, IBB, RanGTP, and FG-Nup binding sites of the importins, as well as the known NES-cargo, RanGTP, and FG-Nup binding sites of the exportins. These binding sites are obtained from the crystal structures of the bound states of Kapβs using the PiSITE webserver [52], giving results for the importins KAP121, Impβ1, TNPO1, KAP95, and TNPO3, and the exportins KAP124, CRM1, and XPO5. The crystal structures used to find the binding sites are listed in table S3 of the SI. The A- and B-helices are also highlighted in figure 3a. As expected, for all cases an increasing poly-PR length increases the contact probability for individual residues. More interestingly, we observe that, depending on the poly-PR length and the type of Kapβ, poly-PR can make contact with Kapβ residues that are also used for the recognition of cargo, IBB domain, RanGTP, and FG-Nups. In figure 3b we show the number of contact residues shared between poly-PR and the native binding partners of the Kapβs, *N*_shared_. It is seen that poly-PR makes contacts with importins at several known cargo-NLS, IBB, and RanGTP binding residues. At a poly-PR-importin molar ratio of 1:1, we find no overlap between poly-PR and the known FG-Nup binding sites. For all importins and exportins, the number of shared contact sites between poly-PR and the native binding partners of Kapβs increases by increasing poly-PR length (see figure 3b).

**Figure 3:**
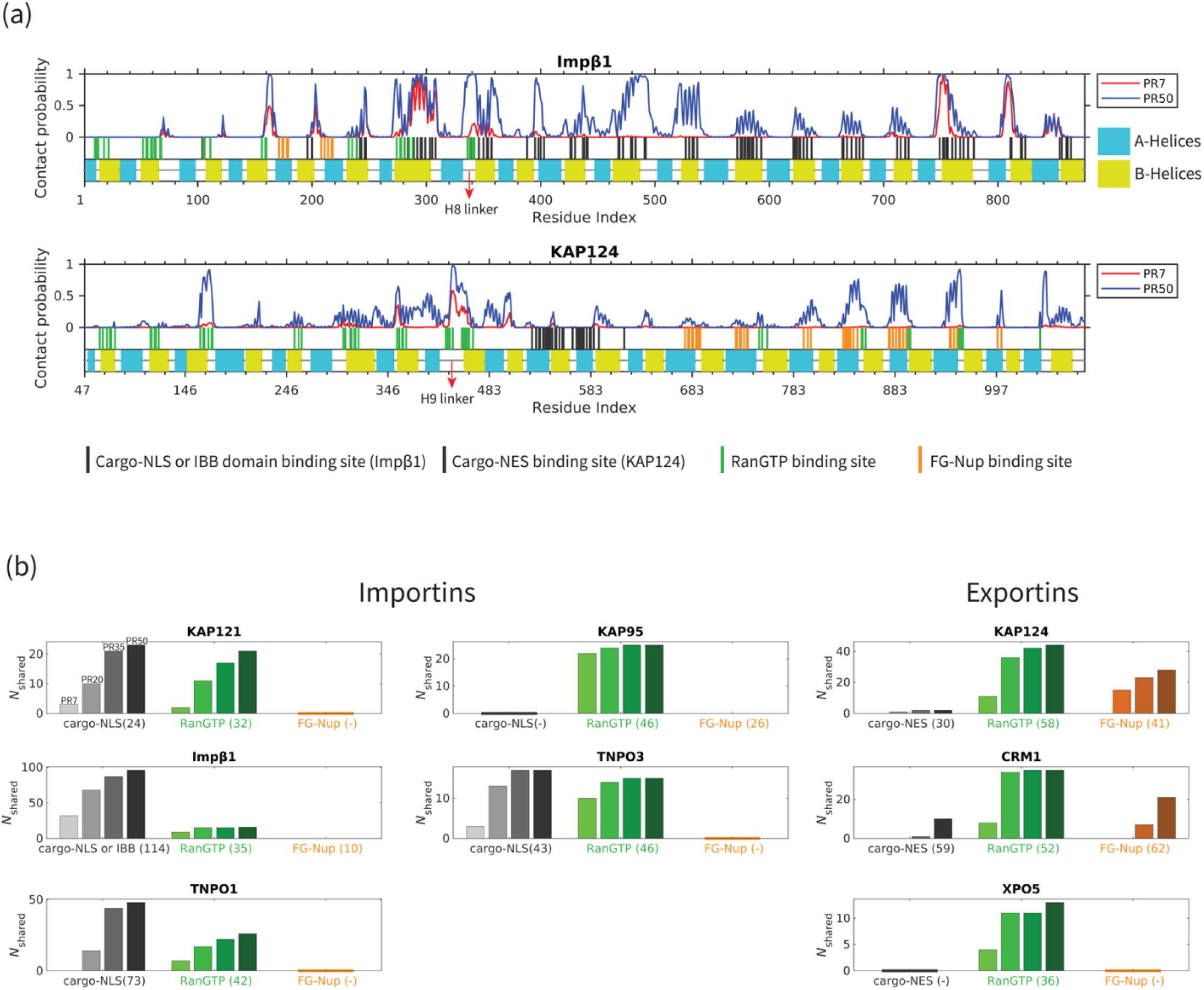
Poly-PR interacts with Kapβs sites used for the recognition of NLS-cargo, IBB, RanGTP and FG-Nups in a poly-PR-length-dependent manner. (a) The contact probability of each Kapβ residue in interacting with poly-PR plotted against the residue index for the Impβ1 importin and the KAP124 exportin at monovalent salt concentration *C*_salt_ = 100 mM. See figure S6 for the results for other Kapβs. In each figure, the curves correspond to PR7 and PR50. In the bottom part of each figure, the first row shows the known binding sites for NLS/NES-cargo, IBB domain, RanGTP, and FG-Nups. These binding sites are obtained from the crystal structures of the bound states of Kapβs in the Protein Data Bank using PiSITE, see table S3 of the SI for more details. For importins, residues that bind to NLS-cargo and IBB domain, and for exportins residues that bind to NES-cargo, are shown with vertical black lines. The residues that bind to RanGTP and FG-Nups are shown with vertical green and orange lines, respectively. The RanGTP data contains binding residues for both RanGTP and RanGppNHp which is the non-hydrolysable form of RanGTP. The second row in each figure shows the A- and B-helices in light blue and yellow, respectively. The linkers that connect the helices are shown with grey horizontal lines. The H8 loop of Impβ1 and the H9 loop of KAP124 are highlighted with red arrows. (b) Number of contact sites shared between poly-PR and the binding partners of Kapβs, *N*_shared_, plotted for PR7, PR20, PR35, and PR50. In each bar plot, the numbers inside the parentheses on the horizontal axis shows the number of the known binding sites obtained from PiSITE. We use a (-) mark if there is no known binding site. As can be seen, for importins, a limited number of FG-Nup binding sites are available. In each set of bar plots, we report the results for PR7, PR20, PR35, and PR50 (from left to right). Bars with darker colors correspond to longer poly-PR chains.

A common feature among the importins is the interaction of poly-PR with the linker between A and B helices of HEAT 8 (H8 linker), see figures 3a and S6. This specific linker region is highly acidic in several importins, including KAP121, Imp5, Impβ1, TNPO1, and KAP95, and has been shown to play a role in binding of RanGTP [53-56], the IBB domain [42], and NLS-cargo [57]. In the case of TNPO1, the long negatively-charged H8 linker has also been shown to play an important role in cargo release upon binding of RanGTP to the importin [58]. For exportins, poly-PR mostly interacts with RanGTP and FG-Nup binding residues, and less with cargo-NES binding sites (see figure 3a and b). For KAP124 and CRM1, poly-PR interacts with a relatively long linker that connects the A and B helices of HEAT 9 (H9 linker) with a high probability. The H9 linker contains negatively-charged residues and interacts with RanGTP, a process which is necessary for cargo-NES loading [59]. In the absence of RanGTP, the H9 loop and the C-terminal alpha-helix inhibit the binding of cargo-NES to the exportin [41].

At higher poly-PR concentrations, more residues of the Kapβs interact with poly-PR, as shown by the increase of *N*_contact_ in figure S7, S8, and the increase of *N*_shared_ in figure S10. The degree of increase depends on the type of the Kapβ and is more pronounced for shorter poly-PRs. Similar to the effect of the poly-PR length on the interaction with the importins, increasing poly-PR concentration also increases the number of poly-PR contact sites on the A-helices of the importins, see figure S9. The outer surface of the Kapβs, comprising the A-helices, is known to interact with FG-Nups [60-63]. Although for importins, poly-PR only interacts with a limited number of known FG-Nup binding residues, see figure S10 for KAP95 interacting with poly-PR (number of repeat units ≥ 35), the interaction of poly-PR with A-helices suggest a likely overlap between poly-PR contact sites and FG-Nup binding sites.

To explore the effect of poly-PR length (at a fixed mass concentration) on its interaction with Kapβs, figure S11 shows the results obtained from simulations performed with the same total number of PR repeat units *n*_PR_ (that is, the same PR mass concentration), but with different lengths of poly-PR. For each Kapβ, we compare the number of contact residues *N*_contact_ for five different *n*_PR_ values. At a certain *n*_PR_, we report the results for two different groups; the first group contains several copies (≥ 3) of PR7, whereas the second group contains fewer copies of a longer poly-PR, i.e. PR20, PR35, PR50, such that *n*_PR_ is the same as in the first group. Figure S11 clearly demonstrates that for all Kapβs the longer poly-PRs cause more contacts for a fixed mass concentration. At lower *n*_PR_ values, where there is only one copy of the longer poly-PR, this effect is less pronounced and for a few cases we even observe a higher *N*_contact_ for shorter poly-PRs. In these cases, one copy of the longer poly-PR mostly interacts with one of the favorable regions of the Kapβ while several copies of PR7 are able to make contact with other regions as well. At higher *n*_PR_ values, with more than one copy of the longer poly-PR, the poly-PRs can roam a larger region and we observe a higher *N*_contact_. At a relatively high mass concentration, longer poly-PRs can potentially interfere more with the function of Kapβ compared to shorter chains, possibly contributing to the poly-PR length-dependence of C9orf72 DPR toxicity.

Based on our results, the following mechanistic picture emerges of the interference of poly-PR with the function of Kapβs as regulators of nucleocytoplasmic transport. In the import cycle, illustrated in figure 4a, poly-PR could interfere with the cargo loading of several Kapβs by interacting with the sites used for the recognition of different cargo-NLSes and the IBB domain. The increased interaction of longer poly-PRs at higher poly-PR concentrations with the A-helices also suggests a potential interference of FG-Nup binding to the outer surface of the importins. Inside the nucleus, poly-PR interacts with RanGTP binding sites, and therefore could cause defects in RanGTP-mediated cargo release and the transport of RanGTP-importin complex back to the cytoplasm. In the export cycle (see figure 4b), poly-PR could interfere with the RanGTP-mediated cargo-NES loading by interacting with RanGTP binding sites. Poly-PR could also interfere with the transport of the exportin from the cytoplasm back to the nucleus through interaction with the FG-Nup binding sites. Based on the poly-PR length-dependent interaction with the known binding sites of Kapβs, see figure 3b, we conclude that a longer poly-PR is likely to play a more important role in the proposed nucleocytoplasmic transport defects described in figure 4.

**Figure 4:**
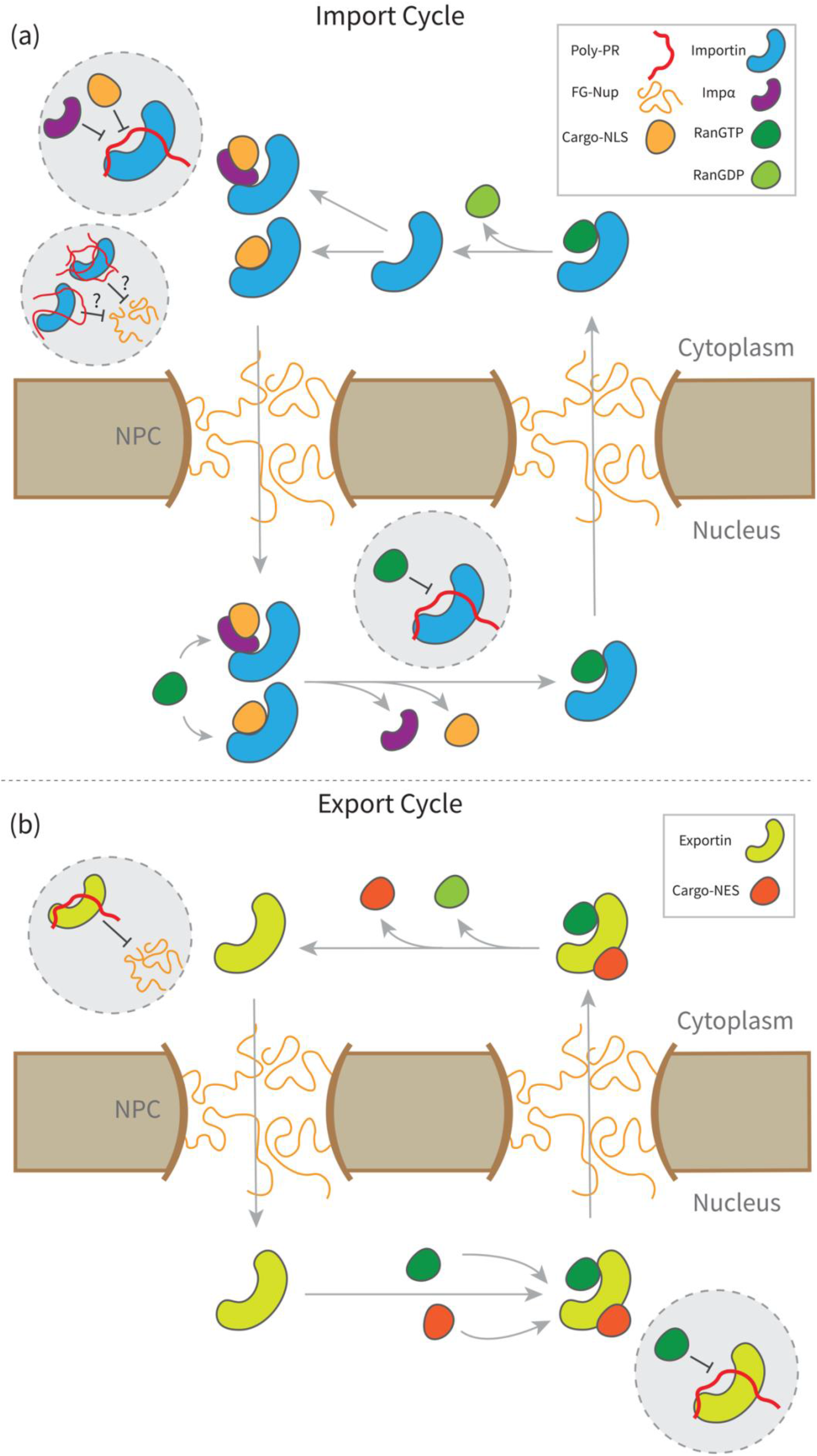
A mechanistic picture for the key molecular interactions that drive poly-PR interference with the function of Kapβs in the nucleocytoplasmic transport. (a) (Top panel) Proposed schematic for the mechanistic pathways of poly-PR interference with the import cycle. Poly-PR could cause defects in cargo loading in the cytoplasm and cargo-release in the nucleus by interaction with NLS-cargo, IBB domain, and RanGTP binding sites of importins. Increasing the length or concentration of poly-PR, increases the interaction of poly-PR with the outer surface of the importin, suggesting a possible change in the way the importin interacts with the FG-Nups. (b) (Bottom panel) Proposed schematic for the mechanistic pathways of poly-PR interference with the export cycle. Poly-PR could cause defects in cargo loading in the nucleus by interaction with the RanGTP binding sites. Poly-PR might also cause defects in the transport of exportin back into the nucleus by interaction with the FG-Nup binding sites.

## Conclusion

We used coarse-grained molecular dynamics models to study the interaction of poly-PR with the unbound state of Kapβs with the aim to gain mechanistic insight in the interference of poly-PR with the functioning of Kapβs in nuclear transport. We showed that poly-PR–Kapβ binding depends on the net charge per residue (NCPR) of the Kapβ, the salt concentration of the solvent, and the poly-PR length. For poly-PR chains with more than 20 repeat units, we observed a linear correlation between the number of poly-PR–Kapβ contacts and the negative NCPR of Kapβs.

We showed that poly-PR tends to make contact with the inner surfaces of most importins (KAP114, KAP121, Imp5, Impβ1, TNPO1, KAP95, and TNPO3) especially at regions with a pronounced negative electrostatic potential. This binding behavior results in the interaction of poly-PR with a large number of cargo-NLS, IBB, and RanGTP binding sites. Our findings also revealed that longer poly-PRs at higher concentrations are able to make contact with the outer surfaces of importins that contain several binding sites for FG-Nups. We also showed that poly-PR binds to exportins, especially at lower salt concentrations, making contacts with several RanGTP and FG-Nup binding sites.

Overall, our results suggest that poly-PR might cause defects in cargo loading, cargo release, and NPC transport of various Kapβs. Furthermore, poly-PR interaction with RanGTP binding sites could also cause defects in the transport of RanGTPs to the cytoplasm, thus interfering with the steep Ran gradient across the nuclear envelop that is necessary to sustain transport. We also observed a pronounced poly-PR length-dependence: increasing poly-PR length increases (i) the poly-PR–Kapβ binding probability, (ii) the contact probability for individual residues of Kapβs, and (iii) the number of contact residues shared between poly-PR and the native binding partners of Kapβs. These findings might provide a molecular basis for the more toxic nature of longer poly-PRs in cell and animal models [8, 11, 44].

In general, the electrostatic-driven binding between poly-PR and Kapβs observed in this study emphasizes that positively-charged poly-PR could interfere with complementary charge-based interactions between Kapβs and their native binding partners. Just how poly-PR and the native binding partners of Kapβs, i.e. cargoes, IBB domains, RanGTP, and FG-Nups compete to bind to the same binding sites on Kapβs would be a very interesting subject for further research.

## Methods

### Coarse-grained model

We use a modified version of the implicit solvent, one-bead-per-amino acid (1BPA) model for disordered proteins developed and applied earlier [25, 33-35, 37, 39, 64]. The bonded potentials in this force field are residue specific and sequence specific, and depend on the patterning of three groups of amino acids, i.e. G, P, and other residues. For non-bonded poly-PR-poly-PR interactions, we take into account hydrophobic/hydrophilic and electrostatic interactions. The interactions between poly-PR and Kapβs can be classified in three categories: electrostatic interactions, cation-pi interactions, and excluded volume interactions. The poly-PR model used in this study has been applied earlier to study the phase separation of DPRs [37]. For Kapβs, a network of stiff harmonic bonds is used to maintain the secondary and tertiary structure of the protein. The missing regions in the crystal structures of the Kapβs have more than 50% of their residues in the coil conformation, as shown in figure S1 based on the results from the PSIPRED predictor [65], and are included in the CG model as disordered regions. A more in-depth discussion on the force field is provided in section 1 of the SI.

### Simulation and contact analysis

Langevin dynamics simulations are performed at 300 K at monovalent salt concentrations of 100 mM and 200 mM in NVT ensembles with a time-step of 0.02 ps and a Langevin friction coefficient of 0.02 ps^-1^ using GROMACS version 2018. Simulations are performed for at least 2.5 µs in cubic periodic boxes, and the last 2 µs are used for the analyzing the interaction between poly-PR and the Kapβs. To calculate the number of contacts, binding probability, and contact probability for individual residues, a cut-off of 1 nm is used. The length of A- and B-helices are estimated using the STRIDE algorithm in VMD [47], and depicted using the Bendix plugin in VMD [66, 67]. The binding sites are obtained from the crystal structures of the bound states of Kapβs in the Protein Data Bank using PiSITE [52]. The electrostatic potentials are calculated using PDB2PQR [51], and are shown using Surf representation in VMD on a red-white-blue map. Positive and negative surface potentials are drawn in blue and red. Additional details are provided in sections 2-4 of the SI.

## References

1. DeJesus-Hernandez, M., et al., Expanded GGGGCC Hexanucleotide Repeat in Noncoding Region of C9ORF72 Causes Chromosome 9p-Linked FTD and ALS. Neuron, 2011. 72(2): p. 245–256.

2. Renton, A.E., et al., A hexanucleotide repeat expansion in C9ORF72 is the cause of chromosome 9p21-linked ALS-FTD. Neuron, 2011. 72(2): p. 257–268.

3. Rohrer, J.D., et al., C9orf72 expansions in frontotemporal dementia and amyotrophic lateral sclerosis. The Lancet Neurology, 2015. 14(3): p. 291–301.

4. Mori, K., et al., The C9orf72 GGGGCC repeat is translated into aggregating dipeptide-repeat proteins in FTLD/ALS. Science, 2013. 339(6125): p. 1335–1338.

5. Zu, T., et al., RAN proteins and RNA foci from antisense transcripts in C9ORF72 ALS and frontotemporal dementia. Proceedings of the National Academy of Sciences of the United States of America, 2013. 110(51): p. E4968–E4977.

6. Taylor, J.P., R.H. Brown, and D.W. Cleveland, Decoding ALS: From genes to mechanism. Nature, 2016. 539(7628): p. 197–206.

7. Gendron, T.F., et al., Antisense transcripts of the expanded C9ORF72 hexanucleotide repeat form nuclear RNA foci and undergo repeat-associated non-ATG translation in c9FTD/ALS. Acta Neuropathologica, 2013. 126(6): p. 829–844.

8. Mizielinska, S., et al., C9orf72 repeat expansions cause neurodegeneration in Drosophila through arginine-rich proteins. Science, 2014. 345(6201): p. 1192–1194.

9. Belzil, V.V., et al., Reduced C9orf72 gene expression in c9FTD/ALS is caused by histone trimethylation, an epigenetic event detectable in blood. Acta Neuropathologica, 2013. 126(6): p. 895–905.

10. Kwon, I., et al., Poly-dipeptides encoded by the C9orf72 repeats bind nucleoli, impede RNA biogenesis, and kill cells. Science, 2014. 345(6201): p. 1139–1145.

11. Jovicic, A., et al., Modifiers of C9orf72 dipeptide repeat toxicity connect nucleocytoplasmic transport defects to FTD/ALS. Nature Neuroscience, 2015. 18(9): p. 1226–1229.

12. Wen, X., et al., Antisense proline-arginine RAN dipeptides linked to C9ORF72-ALS/FTD form toxic nuclear aggregates that initiate invitro and invivo neuronal death. Neuron, 2014. 84(6): p. 1213–1225.

13. Lee, K.H., et al., C9orf72 Dipeptide Repeats Impair the Assembly, Dynamics, and Function of Membrane-Less Organelles. Cell, 2016. 167(3): p. 774-788.e17.

14. Shi, K.Y., et al., Toxic PRn poly-dipeptides encoded by the C9orf72 repeat expansion block nuclear import and export. Proceedings of the National Academy of Sciences of the United States of America, 2017. 114(7): p. E1111–E1117.

15. Zhang, Y.J., et al., Heterochromatin anomalies and double-stranded RNA accumulation underlie C9orf72 poly(PR) toxicity. Science, 2019. 363(6428).

16. Boeynaems, S., et al., Drosophila screen connects nuclear transport genes to DPR pathology in c9ALS/FTD. Scientific Reports, 2016. 6(1): p. 20877–20877.

17. Balendra, R. and A.M. Isaacs, C9orf72-mediated ALS and FTD: multiple pathways to disease. Nature Reviews Neurology, 2018. 14(9): p. 544–558.

18. Hutten, S. and D. Dormann, Nucleocytoplasmic transport defects in neurodegeneration - Cause or consequence? Semin Cell Dev Biol, 2020. 99: p. 151–162.

19. Hutten, S., et al., Nuclear Import Receptors Directly Bind to Arginine-Rich Dipeptide Repeat Proteins and Suppress Their Pathological Interactions. Cell Rep, 2020. 33(12): p. 108538.

20. Hayes, L.R., L. Duan, K. Bowen, P. Kalab, and J.D.J.E. Rothstein, C9orf72 arginine-rich dipeptide repeat proteins disrupt karyopherin-mediated nuclear import. eLife, 2020. 9: p. e51685.

21. Stewart, M., Molecular mechanism of the nuclear protein import cycle. Nat Rev Mol Cell Biol, 2007. 8(3): p. 195–208.

22. Strambio-De-Castillia, C., M. Niepel, and M.P. Rout, The nuclear pore complex: bridging nuclear transport and gene regulation. Nat Rev Mol Cell Biol, 2010. 11(7): p. 490–501.

23. Timney, B.L., et al., Simple rules for passive diffusion through the nuclear pore complex. Journal of Cell Biology, 2016. 215(1).

24. Mohr, D., S. Frey, T. Fischer, T. Güttler, and D. Görlich, Characterisation of the passive permeability barrier of nuclear pore complexes. Embo j, 2009. 28(17): p. 2541–53.

25. Popken, P., A. Ghavami, P.R. Onck, B. Poolman, and L.M. Veenhoff, Size-dependent leak of soluble and membrane proteins through the yeast nuclear pore complex. Molecular Biology of the Cell, 2015. 26(7): p. 1386–1394.

26. O’Reilly, A.J., J.B. Dacks, and M.C. Field, Evolution of the karyopherin-β family of nucleocytoplasmic transport factors; ancient origins and continued specialization. PLoS One, 2011. 6(4): p. e19308.

27. Pumroy, R.A. and G. Cingolani, Diversification of importin-α isoforms in cellular trafficking and. Biochemical Journal, 2015. 466: p. 13–28.

28. Mitrousis, G., A.S. Olia, N. Walker-Kopp, and G. Cingolani, Molecular basis for the recognition of snurportin 1 by importin. Journal of Biological Chemistry, 2008. 283(12): p. 7877–7884.

29. Lott, K. and G. Cingolani, The importin β binding domain as a master regulator of nucleocytoplasmic transport. Biochim Biophys Acta, 2011. 1813(9): p. 1578–92.

30. Kalita, J., L.E. Kapinos, and R.Y.H. Lim, On the asymmetric partitioning of nucleocytoplasmic transport - recent insights and open questions. J Cell Sci, 2021. 134(7).

31. Görlich, D., M.J. Seewald, and K. Ribbeck, Characterization of Ran-driven cargo transport and the RanGTPase system by kinetic measurements and computer simulation. Embo j, 2003. 22(5): p. 1088–100.

32. Boeynaems, S., et al., Phase Separation of C9orf72 Dipeptide Repeats Perturbs Stress Granule Dynamics. Molecular Cell, 2017. 65(6): p. 1044-1055.e5.

33. Ghavami, A., L.M. Veenhoff, E. Van der Giessen, and P.R. Onck, Probing the disordered domain of the nuclear pore complex through coarse-grained molecular dynamics simulations. Biophysical Journal, 2014. 107(6): p. 1393–1402.

34. Ananth, A.N., et al., Spatial structure of disordered proteins dictates conductance and selectivity in nuclear pore complex mimics. eLife, 2018. 7: p. 1–24.

35. Jafarinia, H., E. Van der Giessen, and P.R. Onck, Phase Separation of Toxic Dipeptide Repeat Proteins Related to C9orf72 ALS/FTD. Biophys J, 2020. 119(4): p. 843–851.

36. Fragasso, A., et al., A designer FG-Nup that reconstitutes the selective transport barrier of the nuclear pore complex. Nat Commun, 2021. 12(1): p. 2010.

37. Ketterer, P., et al., DNA origami scaffold for studying intrinsically disordered proteins of the nuclear pore complex. Nature Communications, 2018. 9(1): p. 1–8.

38. Bernardes, N.E. and Y.M. Chook, Nuclear import of histones. Biochemical Society Transactions, 2020. 48(6): p. 2753–2767.

39. Ghavami, A., E. Van der Giessen, and P.R. Onck, Coarse-Grained Potentials for Local Interactions in Unfolded Proteins. Journal of Chemical Theory and Computation, 2013. 9(1): p. 432–440.

40. Yamazawa, R., et al., Structural Basis for Selective Binding of Export Cargoes by Exportin-5. Structure, 2018. 26(10): p. 1393-1398.e2.

41. Saito, N. and Y. Matsuura, A 2.1-Å-resolution crystal structure of unliganded CRM1 reveals the mechanism of autoinhibition. J Mol Biol, 2013. 425(2): p. 350–64.

42. Cingolani, G., C. Petosa, K. Weis, and C.W. Müller, Structure of importin-β bound to the IBB domain of importin-α. Nature, 1999. 399(6733): p. 221–229.

43. Freibaum, B.D., et al., GGGGCC repeat expansion in C9orf72 compromises nucleocytoplasmic transport. Nature, 2015. 525(7567): p. 129–133.

44. Kanekura, K., et al., Characterization of membrane penetration and cytotoxicity of C9orf72-encoding arginine-rich dipeptides. Scientific Reports, 2018. 8(1).

45. White, M.R., et al., C9orf72 Poly(PR) Dipeptide Repeats Disturb Biomolecular Phase Separation and Disrupt Nucleolar Function. Molecular Cell, 2019. 74(4): p. 713-728.e6.

46. Colwell, L.J., M.P. Brenner, and K. Ribbeck, Charge as a selection criterion for translocation through the nuclear pore complex. PLoS Comput Biol, 2010. 6(4): p. e1000747.

47. Frishman, D. and P. Argos, Knowledge-based protein secondary structure assignment. Proteins, 1995. 23(4): p. 566–79.

48. Nanaura, H., et al., C9orf72-derived arginine-rich poly-dipeptides impede phase modifiers. Nat Commun, 2021. 12(1): p. 5301.

49. Okada, C., et al., A high-resolution structure of the pre-microRNA nuclear export machinery. Science, 2009. 326(5957): p. 1275–9.

50. Semmelink, M.F.W., et al., Nuclear transport under stress phenocopies transport defects in models of C9Orf72 ALS. bioRxiv, 2022: p. 2022.04.13.488135.

51. Jurrus, E., et al., Improvements to the APBS biomolecular solvation software suite. Protein Sci, 2018. 27(1): p. 112–128.

52. Higurashi, M., T. Ishida, and K. Kinoshita, PiSite: a database of protein interaction sites using multiple binding states in the PDB. Nucleic Acids Res, 2009. 37(Database issue): p. D360–4.

53. Chook, Y.M. and G. Blobel, Structure of the nuclear transport complex karyopherin-beta2-Ran x GppNHp. Nature, 1999. 399(6733): p. 230–7.

54. Lee, S.J., Y. Matsuura, S.M. Liu, and M. Stewart, Structural basis for nuclear import complex dissociation by RanGTP. Nature, 2005. 435(7042): p. 693–6.

55. Vetter, I.R., A. Arndt, U. Kutay, D. Görlich, and A. Wittinghofer, Structural view of the Ran-Importin beta interaction at 2.3 A resolution. Cell, 1999. 97(5): p. 635–46.

56. Liao, C.C., et al., Karyopherin Kap114p-mediated trans-repression controls ribosomal gene expression under saline stress. EMBO Rep, 2020. 21(7): p. e48324.

57. Kobayashi, J. and Y. Matsuura, Structural basis for cell-cycle-dependent nuclear import mediated by the karyopherin Kap121p. J Mol Biol, 2013. 425(11): p. 1852–1868.

58. Cansizoglu, A.E. and Y.M. Chook, Conformational heterogeneity of karyopherin beta2 is segmental. Structure, 2007. 15(11): p. 1431–41.

59. Monecke, T., et al., Crystal structure of the nuclear export receptor CRM1 in complex with Snurportin1 and RanGTP. Science, 2009. 324(5930): p. 1087–91.

60. Bayliss, R., T. Littlewood, and M. Stewart, Structural basis for the interaction between FxFG nucleoporin repeats and importin-β in nuclear trafficking. Cell, 2000. 102(1): p. 99–108.

61. Liu, S.M. and M. Stewart, Structural basis for the high-affinity binding of nucleoporin Nup1p to the Saccharomyces cerevisiae importin-beta homologue, Kap95p. J Mol Biol, 2005. 349(3): p. 515–25.

62. Bayliss, R., T. Littlewood, L.A. Strawn, S.R. Wente, and M. Stewart, GLFG and FxFG nucleoporins bind to overlapping sites on importin-beta. J Biol Chem, 2002. 277(52): p. 50597–606.

63. Port, S.A., et al., Structural and Functional Characterization of CRM1-Nup214 Interactions Reveals Multiple FG-Binding Sites Involved in Nuclear Export. Cell Rep, 2015. 13(4): p. 690–702.

64. Ghavami, A., E. Van der Giessen, and P.R. Onck, Energetics of Transport through the Nuclear Pore Complex. PLOS ONE, 2016. 11(2): p. e0148876–e0148876.

65. McGuffin, L.J., K. Bryson, and D.T. Jones, The PSIPRED protein structure prediction server. Bioinformatics, 2000. 16(4): p. 404–5.

66. Dahl, A.C., M. Chavent, and M.S. Sansom, Bendix: intuitive helix geometry analysis and abstraction. Bioinformatics, 2012. 28(16): p. 2193–4.

67. Humphrey, W., A. Dalke, and K. Schulten, VMD: visual molecular dynamics. J Mol Graph, 1996. 14(1): p. 33-8, 27-8.

