## Supplementary Information for "Molecular basis of C9orf72 poly-PR interference with the β-karyopherin family of nuclear transport receptors"

### 1 Coarse-grained 1-BPA force field

#### 1.1 Poly-PR–poly-PR interaction

We use the 1-bead-per-amino-acid (1BPA) force field [1, 2] for poly-PR-poly-PR interactions. The 1BPA force field has been previously used to study intrinsically disordered FG-Nups and dipeptide repeat proteins (DPRs) [3, 4]. The bonded interactions, i.e. the bending and torsion potentials, in this force field are residue and sequence specific. The attractive hydrophobic and repulsive hydrophilic interactions between different residues in this force field are represented by:

$$\phi_{\text{hp}} = \begin{cases} \varepsilon_{\text{rep}}\left(\frac{\sigma}{r}\right)^8 - \varepsilon_{ij}\left[\frac{4}{3}\left(\frac{\sigma}{r}\right)^6 - \frac{1}{3}\right] & r \leq \sigma \\ (\varepsilon_{\text{rep}} - \varepsilon_{ij})\left(\frac{\sigma}{r}\right)^8 & r \geq \sigma, \end{cases}$$

where $\varepsilon_{ij} = \varepsilon_{\text{hp}}\sqrt{(\varepsilon_i\varepsilon_j)^{0.27}}$ is the strength of the interaction for each pair of amino acids ($i$,$j$), $r$ is the distance between beads $i$ and $j$, and $\sigma = 0.6$ nm. The values of $\varepsilon_{\text{hp}}$ and $\varepsilon_{\text{rep}}$ are 13 and 10 kJ/mol, respectively. The relative hydrophobic strength values ($\varepsilon_i \in [0,1]$) of the different amino acids are listed in Table S1 [2]. The hydrophobic strength values of charged residues are slightly increased in line with our recent work [4].

The electrostatic interactions between charged residues are described by the modified Coulomb law:

$$\phi_{\text{elec}} = \frac{q_i q_j}{4\pi\varepsilon_0\varepsilon_r(r)r}e^{-\kappa r},$$

where $\varepsilon_r(r) = S_\text{s}\left[1 - \frac{r^2}{z^2}\frac{e^{r/z}}{(e^{r/z}-1)^2}\right]$ is the distance-dependent dielectric constant of the solvent with $S_s = 80$ and $z = 0.25$ nm. The value of the Debye screening coefficient, $\kappa$, is 1 $\text{nm}^{-1}$ for monovalent salt concentration $C_{\text{salt}}$ = 100 mM, and 1.5 $\text{nm}^{-1}$ for $C_{\text{salt}}$ = 200 mM.

#### 1.2 Poly-PR–Kapβ interaction

Poly-PR has been shown to bind to several importins in *in vitro* experiments [5]. However, no binding has been observed for the more hydrophobic DPRs, i.e. poly-GA and poly-GP [5]. These observations suggest the importance of Arginine in driving the binding between poly-PR and the Kapβs. At physiological salt concentrations, Arginine mainly engages in electrostatic and cation-pi interactions. For the poly-PR–Kapβ interaction, we use the same electrostatic potential ($\phi_{\mathrm{elec}}$) as described in the previous section. To take into account the cation-pi interactions between Arginine (in poly-PR) and the aromatic residues Phenylalanine, Tyrosine and Tryptophan (in Kapβ), we use an 8-6 Lennard-Jones (LJ) potential that replaces $\phi_{\mathrm{hp}}$ for the RF, RY, and RW interactions:

$$\phi_{\mathrm{cp},ij}(r) = \varepsilon_{\mathrm{cp},ij}\left[3\left(\frac{r_{\mathrm{m}}}{r}\right)^{8} - 4\left(\frac{r_{\mathrm{m}}}{r}\right)^{6}\right],$$

where $\varepsilon_{\mathrm{cp},ij}$ is a pair-dependent cation-pi energy. The parameter $r_{\mathrm{m}}$, which is the distance at which the $\phi_{\mathrm{cp},ij}$ reaches its minimum value, is set to 0.45 nm. This value is the weighted average distance between the guanidinium group of Arginine and an aromatic ring at different orientations (Planar, Oblique, Orthogonal) [6]. This value of $r_{\mathrm{m}}$ also lies in the range used to find cation-pi structures involving both Arginine and Lysine in the Protein Data Bank (PDB) [7].

The Arginine interaction energy $\varepsilon_{\mathrm{cp},ij}$ with the aromatic side chains of Phenylalanine, Tyrosine, and Tryptophan varies between different pairs [6-9]. In the present study we set the RY cation-pi energy as a basis for calculating the cation-pi energies for the other combinations using PDB statistics. According to all-atom free energy calculations, the RY interaction energy is comparable to the strongest interaction between different non-charged residues at physiological salt concentrations [10]. Therefore, in order for the cation-pi interactions to be compatible with the 1BPA force field, we set $\varepsilon_{\mathrm{cp,RY}} = 5$ kJ/mol which is similar to the deepest potential well in the 1BPA force field (5.2 kJ/mol).

To estimate the energy difference between RY and the other combinations, similar to [11], we use the PDB cation-pi contact frequencies in an aqueous environment [7]. Based on the frequencies of individual residues as well as the frequencies of cation-pi pairs within a large dataset of proteins, see Table S2 taken from [7], the energy differences between different cation-pi pairs can be estimated using a simple formulation of statistical potential [11, 12]. As an example, using $k_{\mathrm{B}}T \approx 2.5$ kJ/mol at $T = 300$ K and $p(\mathrm{F}) = 9162$, $p(\mathrm{Y}) = 8309$, $p(\mathrm{RF}) = 630$, and $p(\mathrm{RY}) = 749$ from Table S2, the energy difference between RY and RF (former minus latter) can be estimated as $-k_{\mathrm{B}}T\ln([p(\mathrm{RY})/p(\mathrm{RF})][p(\mathrm{F})/p(\mathrm{Y})]) \approx -0.7$ kJ/mol. The value of $\varepsilon_{\mathrm{cp,RF}}$ is then $\varepsilon_{\mathrm{cp,RF}} \approx \varepsilon_{\mathrm{cp,RY}} - 0.7$ kJ/mol $= 4.3$ kJ/mol. A similar calculation for the other combinations results in the following cation-pi energies:

| Cation-pi pair | $\varepsilon_{\mathrm{cp,RF}}$ | $\varepsilon_{\mathrm{cp,RY}}$ | $\varepsilon_{\mathrm{cp,RW}}$ | $\varepsilon_{\mathrm{cp,KF}}$ | $\varepsilon_{\mathrm{cp,KY}}$ | $\varepsilon_{\mathrm{cp,KW}}$ |
|---|---|---|---|---|---|---|
| Energy (kJ/mol) | 4.30 | 5.00 | 6.70 | 1.79 | 3.13 | 4.26 |

For the hydrophilic/hydrophobic interactions between poly-PR and the rest of the Kapβ residues (the grey residues in figure 1a), we use $\phi_{\mathrm{hp}}$ with $\varepsilon_{ij} = 10$ kJ/mol which leads to an excluded volume potential that vanishes at $r = 0.6$ nm.

##### 1.3 Developing 1BPA models of Kapβs from the crystal structures

To develop 1BPA coarse-grained (CG) models of the Kapβs we use the crystal structures listed in Table S3. For all the Kapβs listed in this table, except Impβ1 and CRM1, the crystal structure of the unbound state is available. In cases where more than one crystal structure is available, we use the one that has a higher resolution. For Impβ1 (876 residues) and CRM1 (1071 residues), we use Robetta [13] to obtain the crystal structures. Due to the limitation for the sequence length in Robetta, for CRM1, we obtain the structure for residues 72-1071 that includes the C-extension domain which has been shown to play important roles in cargo loading inhibition in the absence of RanGTP [14]. The CG models of Kapβs are built by considering beads at the position of α-carbons in the crystal structures and introducing a network of stiff harmonic bonds that maintains the secondary and tertiary structure of the NTRs. This network of bonds is represented by the harmonic potential $\phi_{\mathrm{network}} = K(r - b)^2$, where $K$ is 8000 kJ/mol/nm$^2$ and $b$ is the original distance between the amino acid beads in the crystal structure. A bond is made between the beads if $b$ is less than 1.4 nm.

There are missing regions in the crystal structure of some Kapβs. Some of these missing regions contain tracts of negatively-charged residues that might play a role in the interaction of the Kapβs with poly-PR. The missing regions in X-ray crystallography are known to be more flexible and more disordered than the observed regions. Here we used PSIPRED to predict the secondary structure of the missing regions [15], see Figure S1. The results show that almost all the missing regions (except a 6-residue-long missing region in a B-helix of KAP121) have more than 50% of their residues in a coil conformation. These regions are added to the CG models and considered to be disordered. For the interactions between the residues within these regions we use the 1BPA force field featuring $\phi_{\mathrm{hp}}$ and $\phi_{\mathrm{elec}}$ as described above.

#### 2 Analyzing the interaction between poly-PR and Kapβs

To analyze the binding between poly-PR and the Kapβs, we calculate the time-averaged number of contacts $C_t$ and the binding probability. For the contact analysis we use a cut-off of 1 nm to find the number of contacts and the binding probabilities in figures 1 and 2. The time-averaged number of contacts between the poly-PR and Kapβ in figures 1 and 2 is obtained by summing the number of contacts per time frame (i.e. the number of poly-PR/Kapβ residue pairs that are within 1 nm) over all frames and dividing by the total number of frames. The binding probability in figures 1 and 2 is the probability of having at least 0.10 of the poly-PR residues within 1 nm proximity of the Kapβ. To calculate the binding probability at equilibrium, we divide the number of frames that satisfy this poly-PR/Kapβ binding criterion by the total number of frames.

The contact probability for each Kapβ residue in figure 3a and S6 is the probability of having at least one poly-PR residue within 1 nm proximity of the Kapβ residue. Similar to the definition of the binding probability, we calculate the contact probability for each Kapβ residue at equilibrium by dividing the number of frames for which this contact criterion is satisfied, by the total number of frames. Residue $i$ is considered to be a contact site if the contact probability for this residue is larger than 0.10. $N_{\text{contact}}$ is the number of Kapβ residues that satisfy this criterion, and $N_{\text{shared}}$ is the number of Kapβ residues that make contact with poly-PR (obtained in our simulations) and at the same time are known for recognition of native binding partners of Kapβs (i.e., NLS/NES-cargo, IBB domain, RanGTP, and FG-Nups, obtained using PiSITE [16], see section 4 for more details).

### 3 Estimating the amino acid sequence of A- and B-helices

To obtain an estimation of the amino acid sequences for the A- and B-helices for each Kapβ, we first use the STRIDE secondary structure prediction algorithm in VMD [17] to find all the α-helices. We then exclude the small helices that usually have smaller than 6-9 residues and consider them to be part of the linkers. In most cases, these small helices are located between two coil regions inside the linkers. However, in a few cases, these helices are connected to larger helices without a coil region in between. For these cases, a helical twist can be seen where the two helices are connected. The approximate location of the twist is found visually and is further checked by obtaining the backbone dihedral angles $\phi$ and $\psi$ for the residues around the twist (see Figure S3). At the location of the twist, the $\psi$ angle changes sign. For Impβ1 and KAP95 we also exclude longer α-helices which contain 13 and 14 residues in the linkers between HEAT repeats 2 and 3. After deleting the α-helices inside the linkers, the number of remaining helices is equal to twice the number of HEAT repeats reported in previous studies, see Table S3 for the information about the HEAT repeats. We exclude KAP120 from our analysis because the number of HEAT repeats has not been reported. We also take into account the exceptions mentioned in the literature, see the comments in Table S4 for Impβ1, TNPO3, and XPO5. A- and B- helices are highlighted in the crystal structures presented in Table S4.

### 4 Using PiSITE to obtain the Kapβ binding sites

We use PiSITE to find residues of Kapβs that interact with protein cargoes, IBB domains, RanGTP, and FG-repeat-containing nucleoporins (FG-Nups). The RanGTP group contains both RanGTP and RanGppNHp (the non-hydrolysable form of RanGTP). This web-based database provides interaction sites of a protein from multiple PDBs including similar proteins. There are a few cargo-importin complexes which are not analyzed by PiSITE. For these cases we find the binding residues based on the information provided in the literature. In this step, only a few new residues (< 5) are added to the list of binding sites which have not been previously predicted by PiSITE. In the last column of Table S3, you can see the list of binding partners for each Kapβ and the corresponding PDBs. Mutated Kapβs are excluded from our analysis.

### 5 Supplementary figures

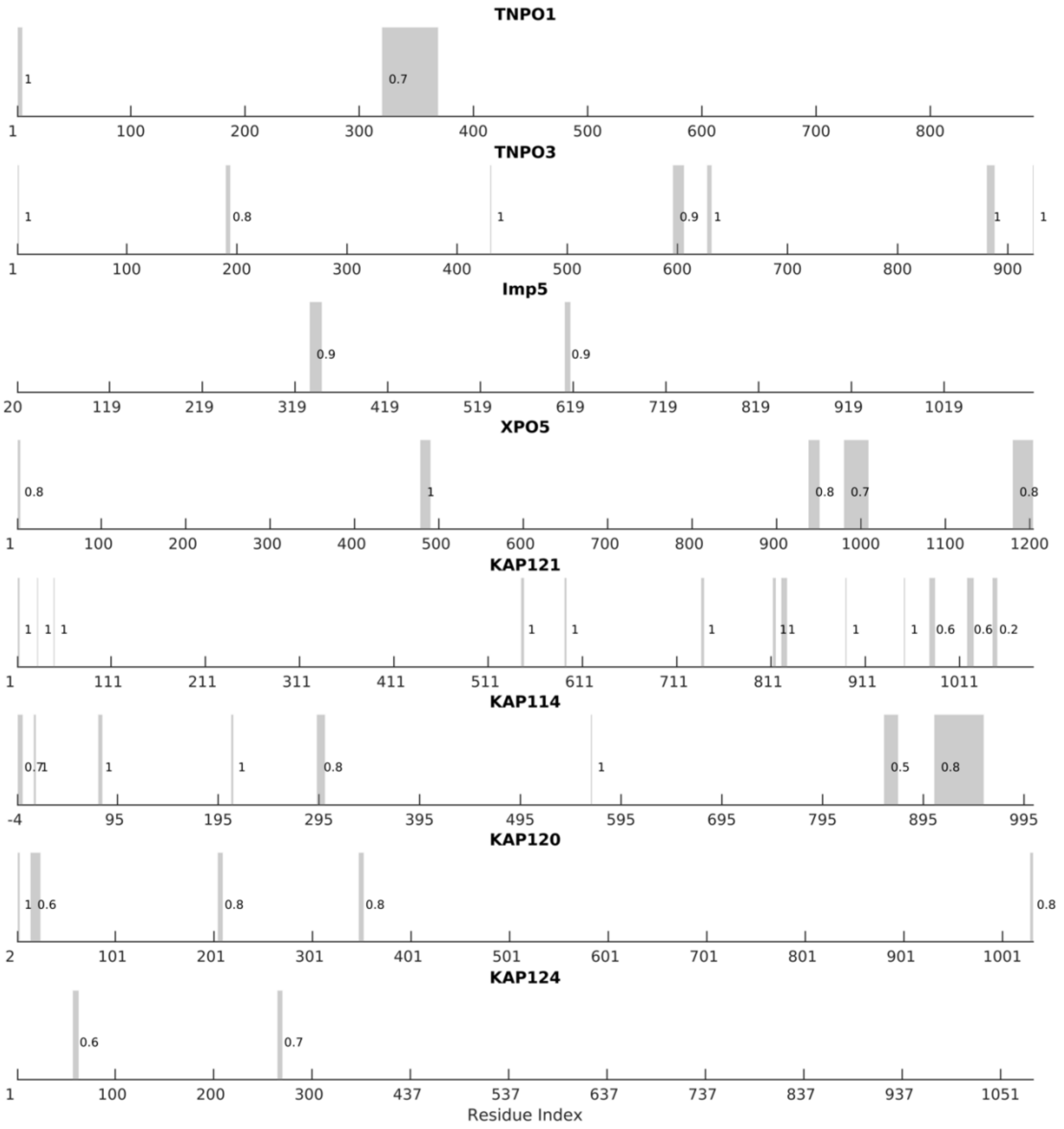

**Figure S1** The missing regions in the structure of different Kapβs (in grey) with the coil conformation probability obtained using PSIPRED depicted next to each region. The coil conformation probability is calculated by dividing the number of residues in a coil conformation by the number of residues in each missing region. All missing regions (except a 6-residue-long region at the C-terminal of KAP121) have a coil conformation probability > 0.50.

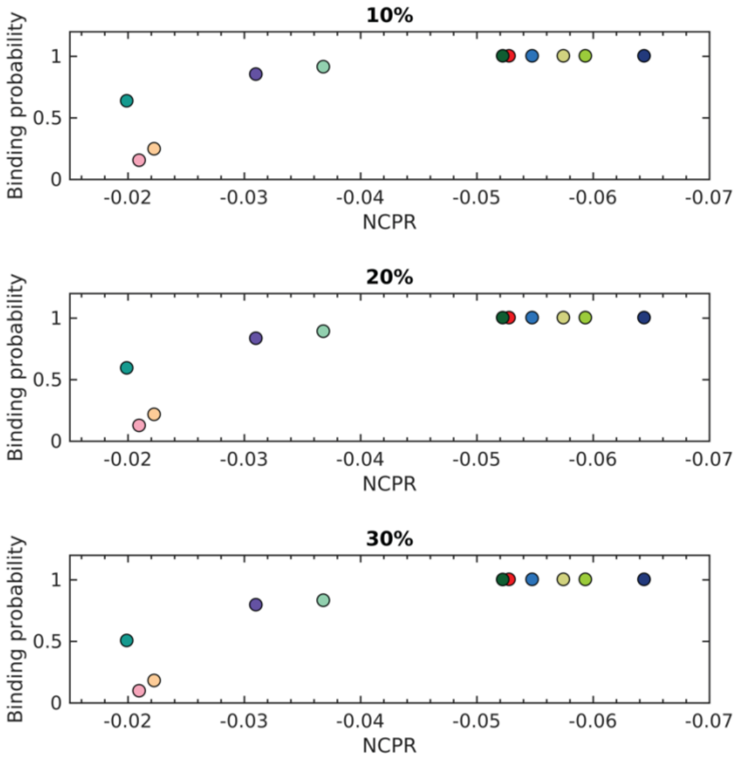

**Figure S2** The results for binding probability $P_{\mathrm{b}}$ using three different values for the binding criterion at $C_{\mathrm{salt}} = 200$ mM. The value on top of each figure shows the threshold for the percentage of poly-PR residues that make contact with Kapβ in the bound state.

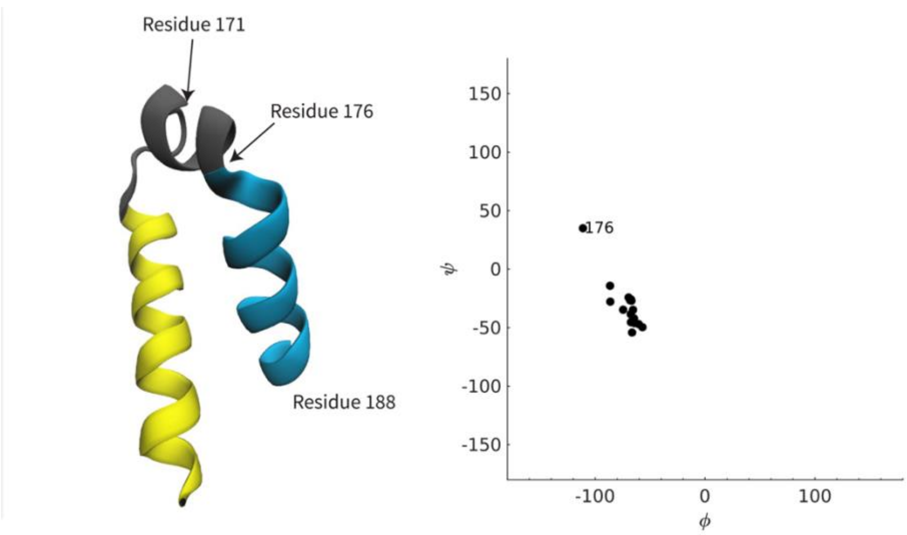

**Figure S3** An example of a local twist in the structure of a Kapβ. (Left panel) Crystal structure of residues 149-188 of KAP95 (PDB code 3nd2). This region contains the B-helix of HEAT 4 (in yellow) and the A-helix of HEAT 5 (in light blue), and the linker region between them (in grey). The location of the local twist is shown with an arrow and is further checked (right panel) by calculating the dihedral angles $\phi$, and $\psi$ for residues 171-188.

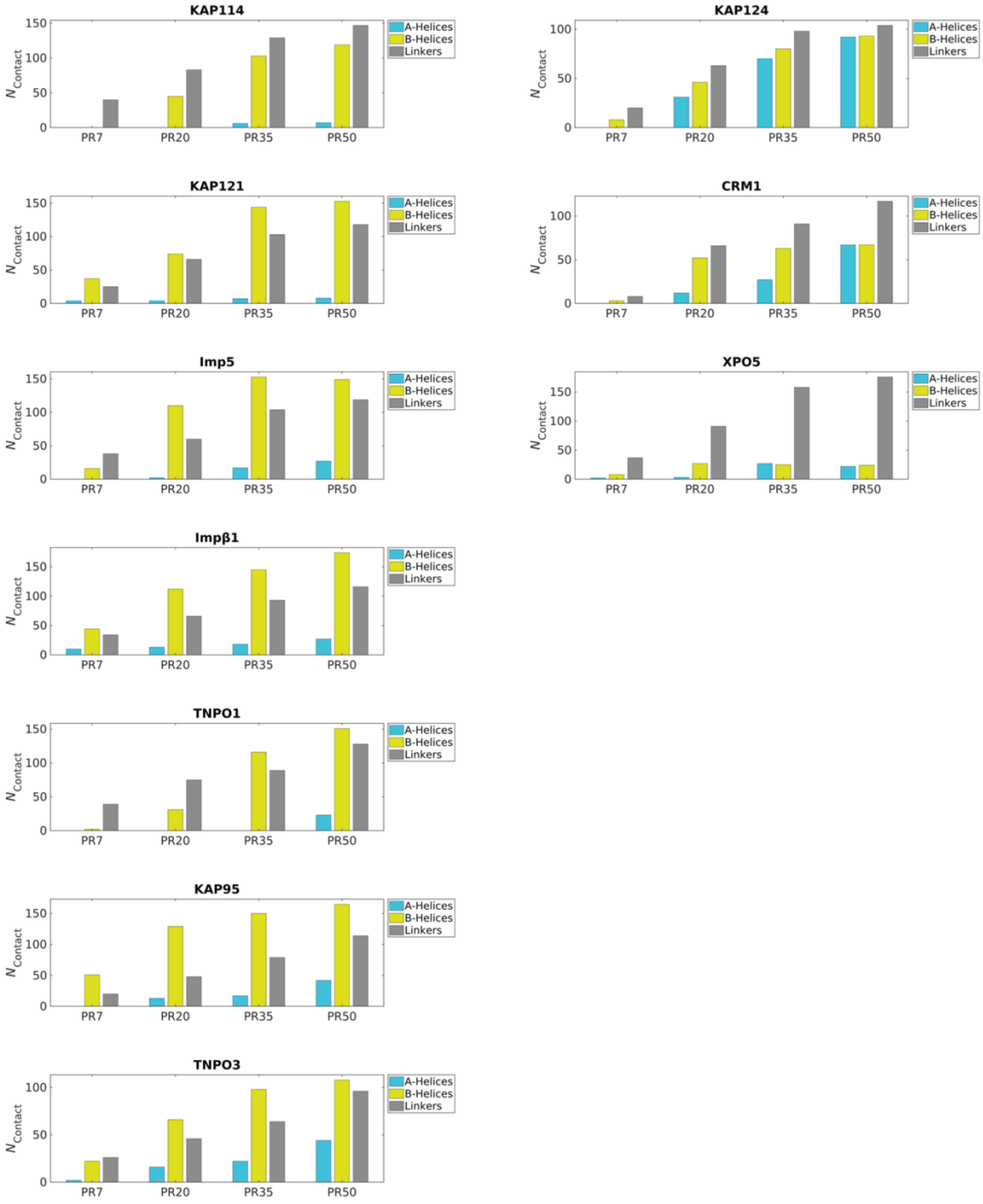

**Figure S4** The number of residues in A-helices, B-helices, and linkers that make contact with poly-PR, $N_{\text{contact}}$, shown for importins (left column) and exportins (right column). The results are reported for PR7, 20, 35, and 50 at $C_{\text{salt}} = 100$ mM.

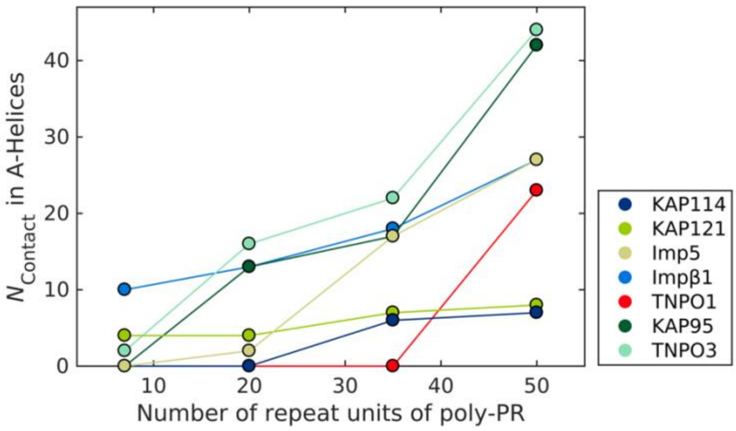

**Figure S5** The number of contact residues in A-helices plotted against poly-PR length for different importins at $C_{\text{salt}} = 100$ mM.

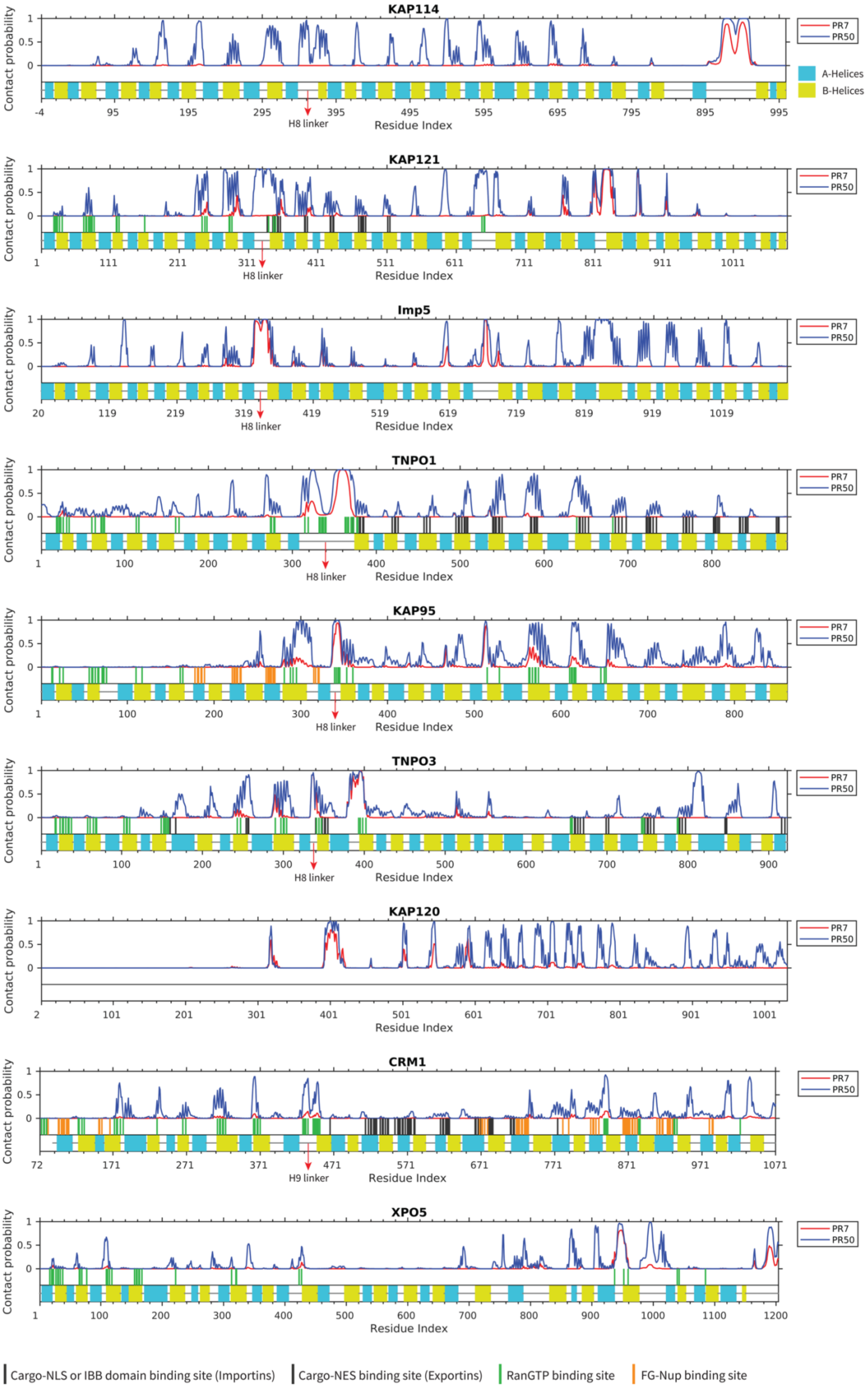
KAP114
KAP121
Imp5
TNPO1
KAP95
TNPO3
KAP120
CRM1
XPO5
Contact probability
Residue Index
PR7
PR50
A-Helices
B-Helices
H8 linker
H9 linker
Cargo-NLS or IBB domain binding site (Importins)
Cargo-NES binding site (Exportins)
RanGTP binding site
FG-Nup binding site

**Figure S6** Contact probability of each Kapβ residue in interacting with PR7 and PR50 plotted against the residue index for different Kapβs at monovalent salt concentration $C_{\mathrm{salt}} = 100$ mM. In the bottom part of each figure, the first row shows the known binding sites for NLS/NES-cargo, IBB domain, RanGTP, and FG-Nups. These binding sites are obtained from the crystal structures of the bound states of Kapβs in the Protein Data Bank using PiSITE, see Table S3 for more details. For importins, residues that bind to NLS-cargo and IBB domains, and for exportins residues that bind to NES-cargo, are shown with vertical black lines. The residues that bind to RanGTP and FG-Nups are shown with vertical green and orange lines respectively. The group RanGTP contains binding residues for both RanGTP and RanGppNHp which is the non-hydrolysable form of RanGTP. The second row in each figure shows the A- and B-helices in light blue and yellow, respectively. The linkers that connect the helices are shown with grey horizontal lines. The H8 loop of importins and the H9 loop of CRM1 is highlighted with red arrows.

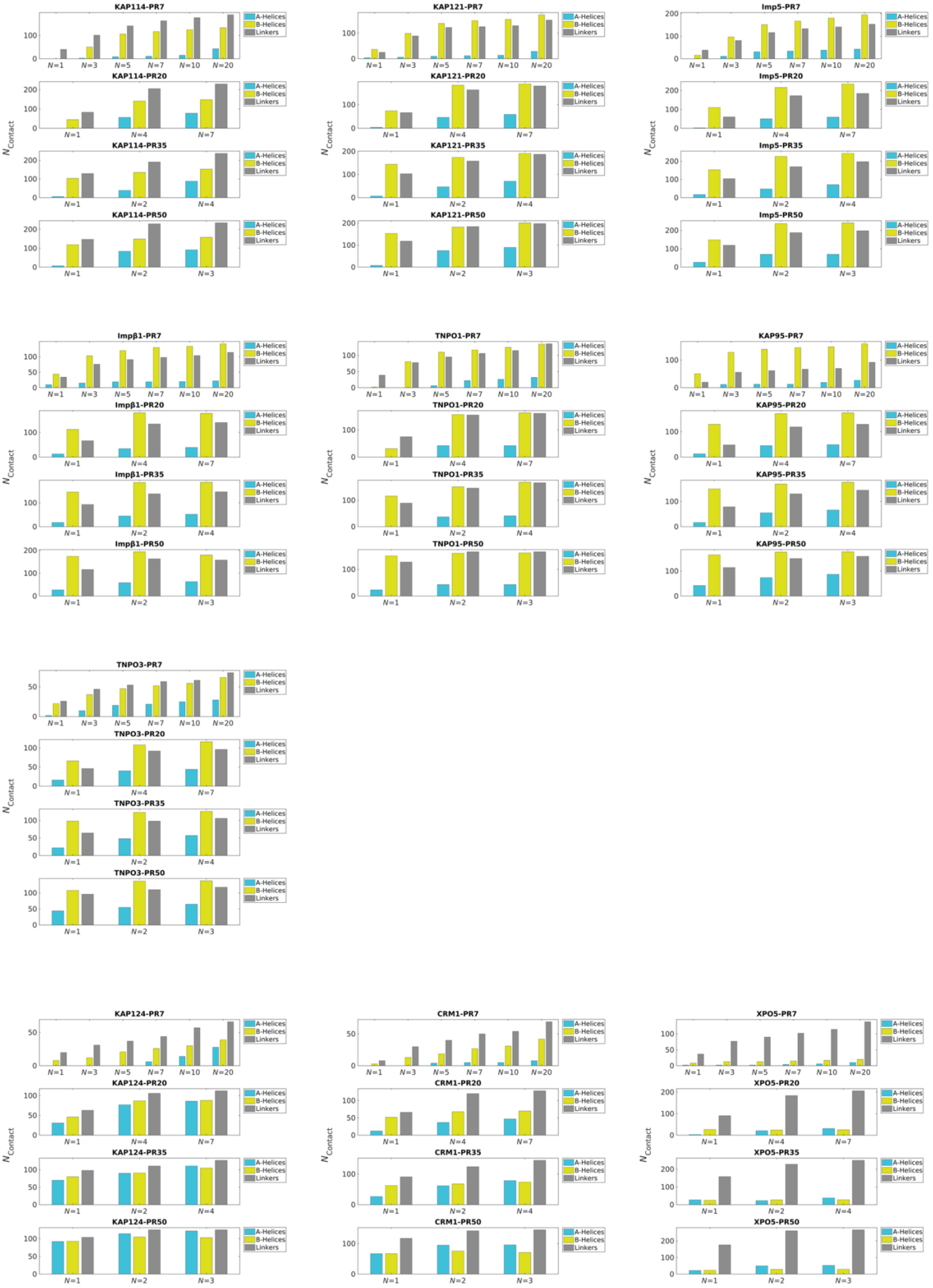

KAP114-PR7
KAP114-PR20
KAP114-PR35
KAP114-PR50
KAP121-PR7
KAP121-PR20
KAP121-PR35
KAP121-PR50
Imp5-PR7
Imp5-PR20
Imp5-PR35
Imp5-PR50
Impβ1-PR7
Impβ1-PR20
Impβ1-PR35
Impβ1-PR50
TNPO1-PR7
TNPO1-PR20
TNPO1-PR35
TNPO1-PR50
KAP95-PR7
KAP95-PR20
KAP95-PR35
KAP95-PR50
TNPO3-PR7
TNPO3-PR20
TNPO3-PR35
TNPO3-PR50
KAP124-PR7
KAP124-PR20
KAP124-PR35
KAP124-PR50
CRM1-PR7
CRM1-PR20
CRM1-PR35
CRM1-PR50
XPO5-PR7
XPO5-PR20
XPO5-PR35
XPO5-PR50
A-Helices
B-Helices
Linkers
N Contact

**Figure S7** The number of residues in each region of the Kapβs that make contact with poly-PR, $N_{\mathrm{contact}}$ at $C_{\mathrm{salt}} = 100$ mM, plotted for different concentrations of PR7, PR20, PR35, and PR50. $N$ is the number of poly-PR molecules.

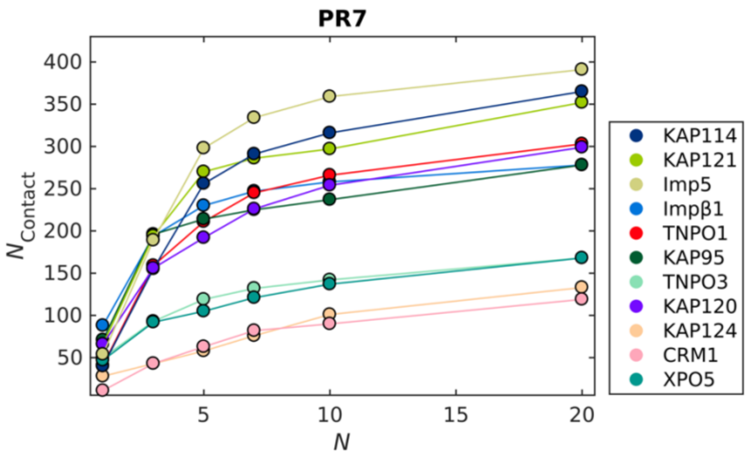

**Figure S8** The number of Kapβ residues that make contact with poly-PR, $N_{\mathrm{contact}}$, at $C_{\mathrm{salt}} = 100$ mM plotted against the number of PR7 molecules, $N$.

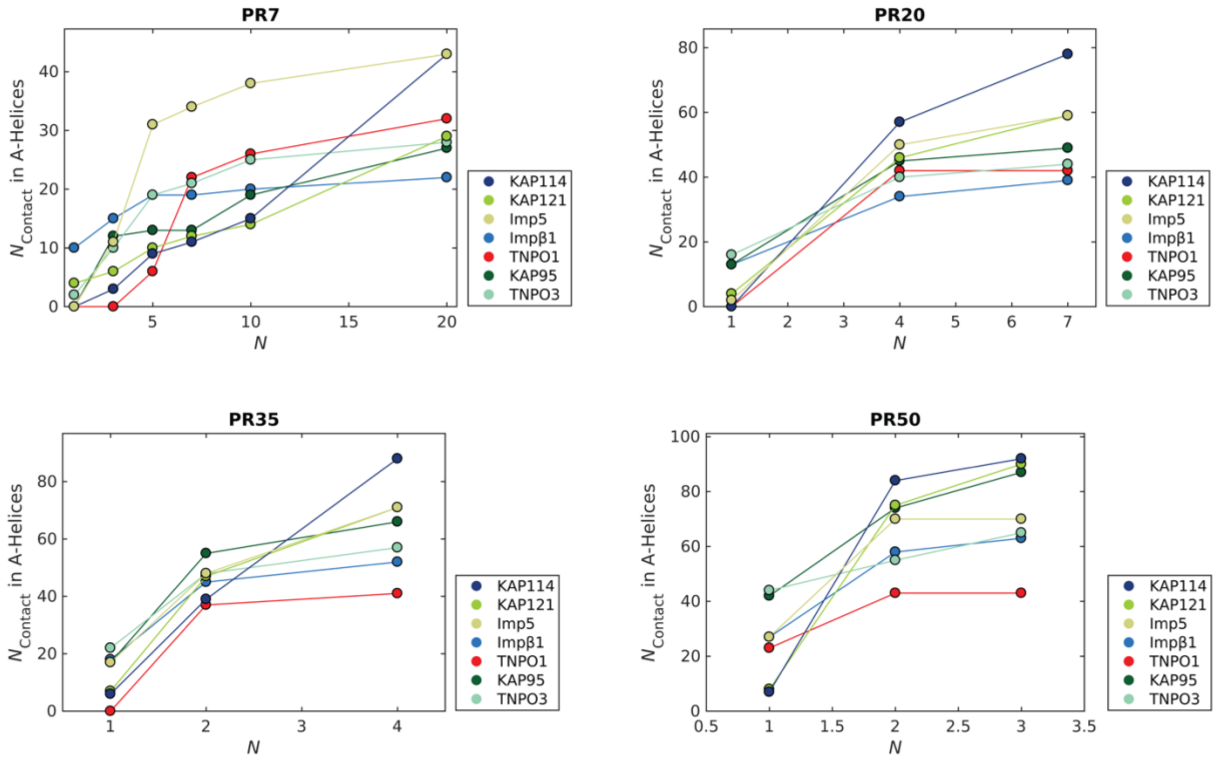

**Figure S9** The number of contact residues in A-helices plotted against the number of poly-PR molecules, $N$, for different importins at $C_{\text{salt}} = 100$ mM.

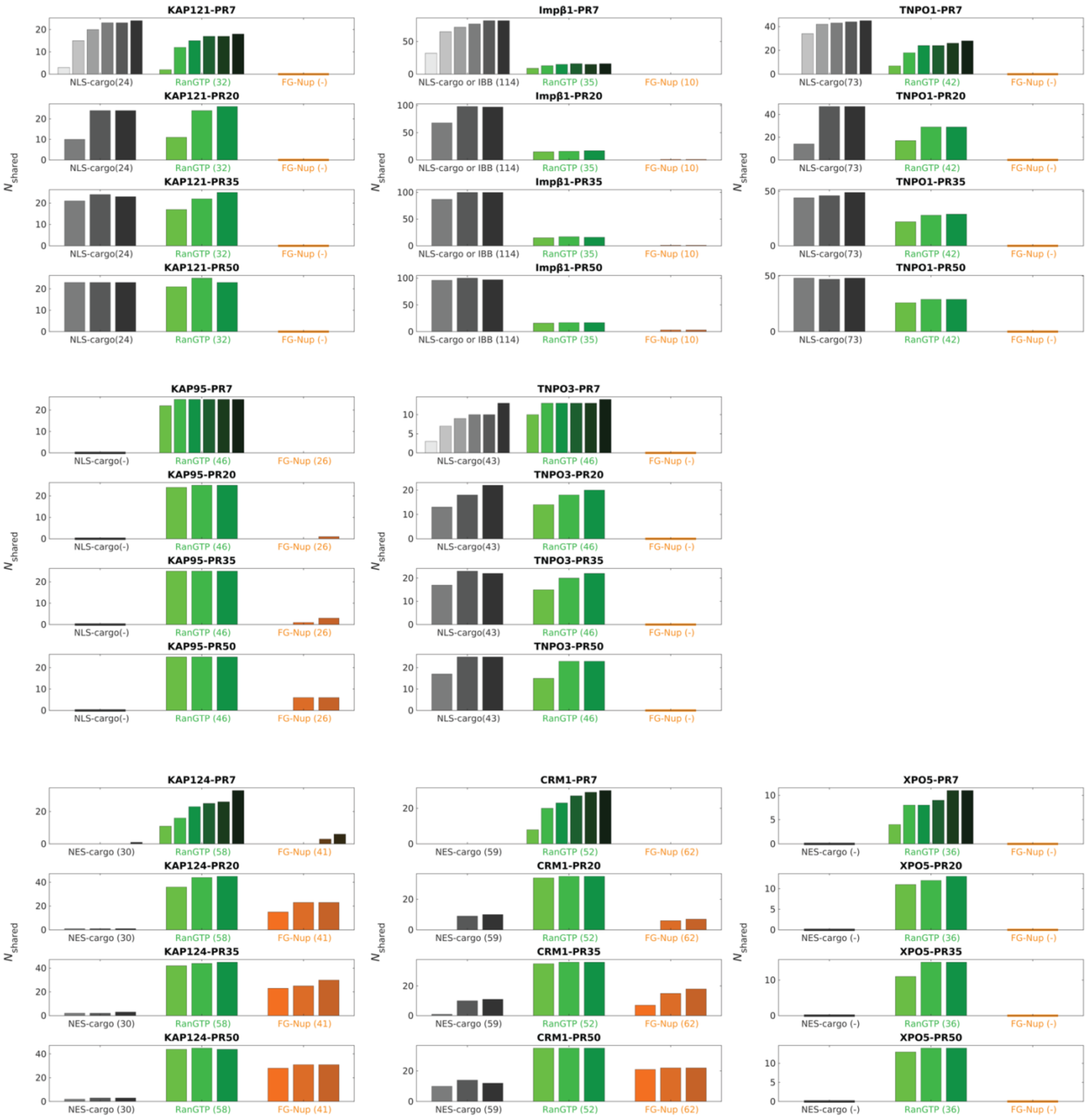

**Figure S10** The number of contact residues shared between poly-PR and the binding partners of Kapβs, $N_{\mathrm{shared}}$, at $C_{\mathrm{salt}} = 100$ mM for different concentrations of poly-PR. The following number of poly-PR molecules, $N$, is used for each poly-PR length. PR7: $N$ = 1, 3, 5, 7, 10, 20. PR20: $N$ = 1, 4, 7. PR35: $N$ = 1, 2, 4. PR50: $N$ = 1, 2, 3. In each set of bar plots, concentration increases from left to right. Bars with darker colors correspond to higher poly-PR concentrations.

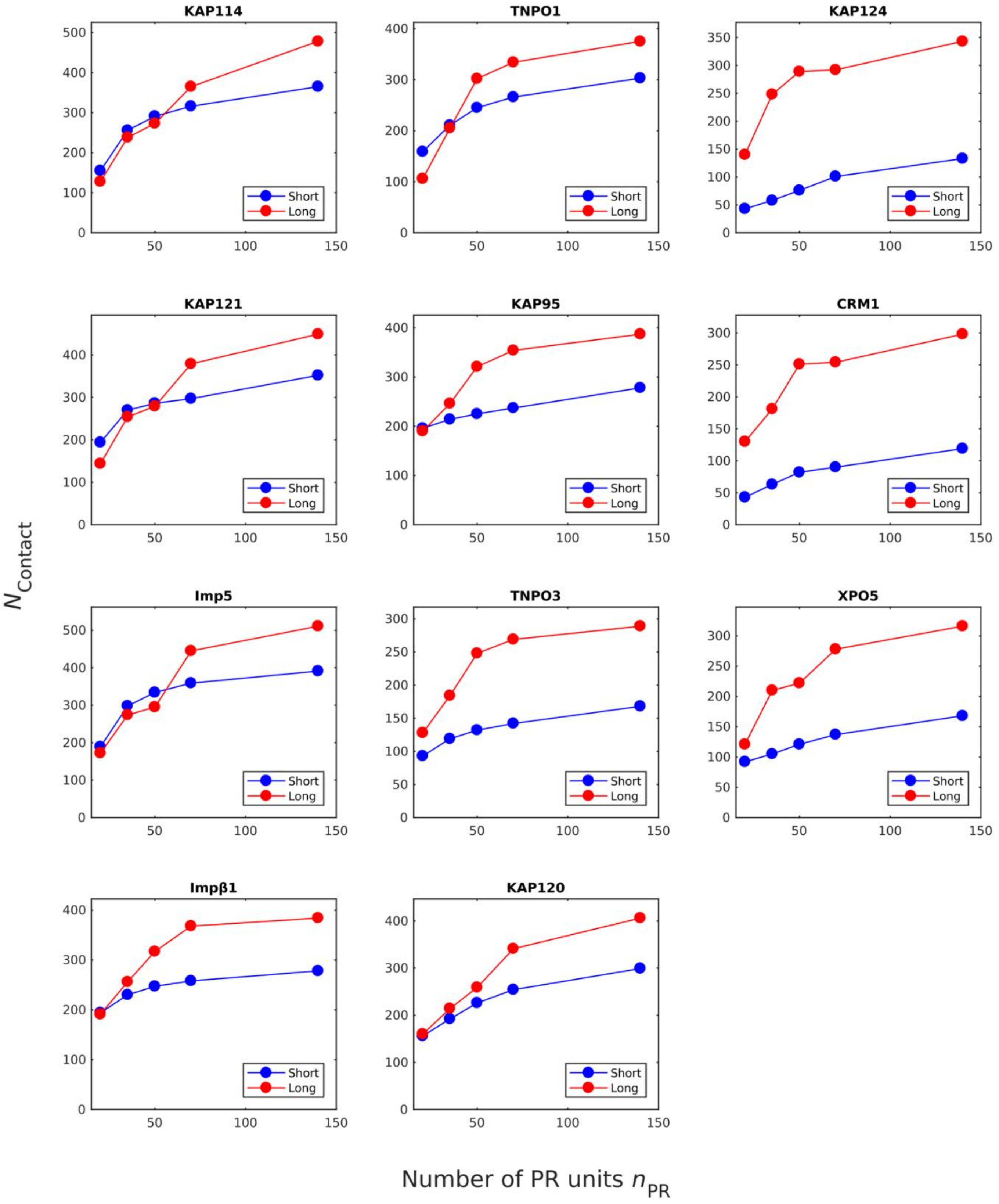

**Figure S11** $N_{\mathrm{contact}}$ plotted against the total number of PR repeat units $n_{\mathrm{PR}}$ in the simulation box for different poly-PR lengths at $C_{\mathrm{salt}} = 100$ mM. At a certain $n_{\mathrm{PR}}$ (or in other words PR mass concentration), the results are reported for two different groups. Group 'short' contains several copies of PR7, whereas the group ‘long’ contains less copies of PR20, 35, or 50. For $n_{\mathrm{PR}}$ = 20 we use (PR7, $N$ =3) and (PR20, $N$ =1), For $n_{\mathrm{PR}}$ = 35 we use (PR7, $N$=5) and (PR35, $N$=1), For $n_{\mathrm{PR}}$ = 50 we use (PR7, $N$=7) and (PR50, $N$=1), For $n_{\mathrm{PR}}$ = 70 we use (PR7, $N$=10) and (PR35, $N$=2). For $n_{\mathrm{PR}}$ = 140 we use PR7, $N$=20) and (PR35, $N$=4).

### 6 Supplementary tables

**Table S1** Relative hydrophobic strength values of the different amino acids [2, 4].

| Amino acid | $\varepsilon_i$ | Amino acid | $\varepsilon_i$ |
|---|---|---|---|
| A | 0.7 | L | 1 |
| R | 0.005 | K | 0.005 |
| N | 0.33 | M | 0.78 |
| D | 0.005 | F | 1 |
| C | 0.68 | P | 0.65 |
| Q | 0.64 | S | 0.45 |
| E | 0.005 | T | 0.51 |
| G | 0.41 | W | 0.96 |
| H | 0.53 | Y | 0.82 |
| I | 0.98 | V | 0.94 |

**Table S2** Frequency of amino acids and cation-pi interactions within proteins (taken from [7]) in a dataset of 593 proteins.

| Amino acid | Total number* | Amino acid pair | Cation-pi interactions** |
|---|---|---|---|
| K | 13446 | KF | 285 |
| R | 10919 | KY | 438 |
| F | 9162 | KW | 283 |
| Y | 8309 | RF | 630 |
| W | 3412 | RY | 749 |
| | | RW | 609 |

*The total number of times a particular amino acid appears in the dataset.
**The number of times a pair of amino acids occurs in a cation-pi interaction.

**Table S3** Information about the Kapβs used in this study.

| **Kapβ** name | Gene name (Uniprot id) | Organism | PDB code | No. of HEATs | HEATs in the CG model (residues) | Binding partners of each Kap**β** (PDB code)** |
|---|---|---|---|---|---|---|
| **Importin subunit beta-1 (Impβ1)** | **KPNB1** (P52292) | Human | Robetta* (Model 1) | 19 | 1-19 (residues 1-876) | IBB domains:<br>Importin-alpha IBB (1qgk), Snurportin-1 IBB (2p8q, 2qna,3lww)<br>Protein cargoes:<br>SREBP-2 (1ukl), SNAIL1(3w5k), PTHrP non-classical NLS (1m5n)<br>Nucleoporins:<br>FXFG repeats from Nsp1p (1f59,1o6o), Synthetic GLFG peptide (1o6p)<br>Ran protein:<br>Ran-GppNHp (1ibr) |
| **Transportin-1 (Kapβ2)** | **TNPO1** (Q92973, isoform 2) | Human | 2qmr | 20 | 1-20 (residues 1-890) | Protein cargoes:<br>hnRNPA1 NLS (2h4m), hnRNPM NLS (2ot8), hnRNPD NLS (2z5n), TAP NLS (2z5k), FUS PY-NLS (4fq3,5yvg,5yvh,5yvi, 4fdd), NAB2 PY-NLS (4jlq), Histone H3 tail (5j3v)<br>Ran protein:<br>Ran-GppNHp (1qbk) |
| **Transportin-3** | **TNPO3** (Q9Y5L0) | Human | 4c0p | 21 | 1-21 (residues 1-923) | Protein cargoes:<br>RNA factor/Splicing factor 2 (ASF/SF2) (4c0o), CPSF6 RSLD (6gx9)<br>Ran protein:<br>Ran-GTP (4ol0) |
| **Importin-5 (Imp5)** | **IPO5** (O00410, isoform 3) | Human | 6xte | 24 | 1-24 (residues 20-1115) | N.A. |
| **Exportin-1 (Exp1)** | **XPO1/CRM1** (O14980) | Human | Robetta* (Model 5) | 20 | 2-20 (residues 72-1071) | Protein cargoes:<br>Snurportin 1 (3gb8,3gjx), PKI NES (3nby), HIV-1 Rev NES (3nbz,3nc0)<br>Ran protein:<br>RanGTP (3gjx,3nby,3nbz,3nc0,3nc1,5dis)<br>Nucleoporins:<br>FG-repeats-containing fragment of Nup214 (5dis) |
| **Exportin-5 (Exp5)** | **XPO5** (Q9HAV4) | Human | 5yu7 | 20 | 1-20 (residues 1-1204) | Ran protein:<br>RanGTP (5yu6) |
| **Importin subunit beta-1** (homolog of human Imp**β**1) | **KAP95** (Q06142) | *S. cerevisiae* (yeast) | 3nd2 | 19 | 1-19 (residues 1-861) | Ran protein:<br>Ran-GTP (2bku) and Ran-GDP (3ea5)<br>Nucleoporins:<br>FG-repeats-containing Nup1p (5owu) |
| **Importin subunit beta-3** (homolog of human IPO5) | **KAP121** (P32337) | *S. cerevisiae* (yeast) | 3w3t | 24 | 1-24 (residues 1-1089) | Protein cargoes:<br>Ste12p (3w3w), Pho4p (3w3x), Cdc14p C-terminus (4zj7), SUMO protease Ulp1p (5h2v)<br>Ran protein:<br>RanGTP (3w3z) |
| **Importin subunit beta-5** (homolog of human Imp9) | **KAP114** (P53067) | *S. cerevisiae* (yeast) | 6aho | 20 | 1-20 (residues 1-1004) | N.A. |
| **Importin beta-like protein KAP120** (homolog of human Imp11) | **KAP120** (Q02932) | *S. cerevisiae* (yeast) | 6fvb | 20 | 1-20 (residues 2-1032) | N.A. |
| **Exportin-1** (homolog of human CRM1) | **KAP124** (P30822) | *S. cerevisiae* (yeast) | 3vyc | 21 | 2-21 (residues 47-1084) | Protein cargoes:<br>PKI NES (3wyg), hRio2 NES (5dhf), CPEB4 NES (5dif), Paxillin NES (5uwh), HDAC5 NES (5uwi), FMRP NES (5uwj), mDia2 NES (5uwp), CDC7 NES (5uwq,5uwr), X11L2 NES (5uws), SMAD4 NES (5uwu)<br>Ran protein:<br>RanGTP (3m1i,3wyf,3wyg,4gmx,4hb2,5dhf,5dif,5uwh, 5uwi,5uwp,5uwq,5uwr,5uws,5uwu)<br>Nucleoporins<br>FG-repeats-containing Nup42 (5xoj) |

* Among the five structures predicted by Robetta, we use the one that has the lowest prediction error.

** For the following cases we add the binding residues suggested in the literature to the list of binding residues obtained from PiSITE. These cases are Impβ1 (PDB code 1m5n [18]), TNPO1 (PDB codes 2z5n [19], 4fdd [20], 5j3v [21]), and TNPO3 (PDB code 6gx9 [22]). These crystal structures are not analyzed by PiSITE. This update only adds a few new binding residues (< 5 residues for each Kapβ) to the list of binding residues obtained from PiSITE.

**Table S4** Our approximation for the A- and B-helices of the Kapβs. For each Kapβ three snapshots are shown: (Left) View down the superhelical or ring-shaped structure of Kapβs, and (middle) side view of the structure of Kapβs with the N-terminal domains on top, and (Right) the electrostatic surface potential of Kapβs. The A- and B-helices are depicted with light blue and yellow tubes using the Bendix plugin in VMD. The electrostatic surface potentials are obtained using PDB2PQR and plotted using the Surf representation in VMD on a red-white-blue map. For better visualization of the electrostatic potential of the inner surfaces of the importins, in some cases the right panel is rotated with respect to the middle panel or part of the electrostatic potential surface is drawn using a transparent surface.

#### Impβ1 (KPNB1)*

| HEAT1 | HEAT2 | HEAT3 | HEAT4 | HEAT5 | HEAT6 | HEAT7 | HEAT8 | HEAT9 | HEAT10 |
|---|---|---|---|---|---|---|---|---|---|
| 3-10,15-31 | 33-45,51-65 | 85-97,108-120 | 128-138,144-160 | 169-181,188-201 | 212-226,231-248 | 260-269,273-303 | 314-331,344-359 | 364-375,380-393 | 399-417,422-438 |
| **HEAT11** | **HEAT12** | **HEAT13** | **HEAT14** | **HEAT15** | **HEAT16** | **HEAT17** | **HEAT18** | **HEAT19** | |
| 448-460,464-486 | 503-514,524-537 | 544-563,571-593 | 599-617,622-639 | 647-660,664-681 | 689-701,710-724 | 731-744,752-777 | 793-806,812-829 | 831-838,841-852,856-874 | |

*In the last helix, there are two A-helices (residues 831-838, and 841-852) connected with a small turn (residues 839, 840) [23]. In our contact site analysis and the visualization presented here, these two residues are considered to be part of the A-helix. Therefore, the last A-helix contains residues 831-852.

*Inside the linker between HEATs 2 and 3, there is a relatively long α-helix with 14 residues [23].

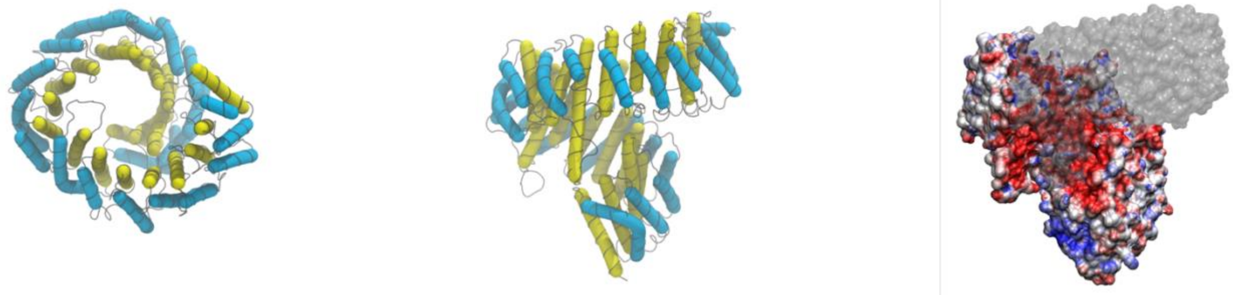

#### TNPO1

| HEAT1 | HEAT2 | HEAT3 | HEAT4 | HEAT5 | HEAT6 | HEAT7 | HEAT8 | HEAT9 | HEAT10 |
|---|---|---|---|---|---|---|---|---|---|
| 7-22,27-38 | 44-54,62-78 | 85-97,104-120 | 128-137,142-158 | 172-183,188-200 | 207-222,229-245 | 253-266,270-285 | 296-307,375-390 | 398-407,411-424 | 435-447,452-471 |
| **HEAT11** | **HEAT12** | **HEAT13** | **HEAT14** | **HEAT15** | **HEAT16** | **HEAT17** | **HEAT18** | **HEAT19** | **HEAT20** |
| 477-489,494-511 | 519-532,535-552 | 559-574,582-597 | 605-628,638-655 | 667-677,681-697 | 706-716,722-739 | 747-759,765-781 | 793-802,808-823 | 832-840,847-864 | 866-875,878-888 |

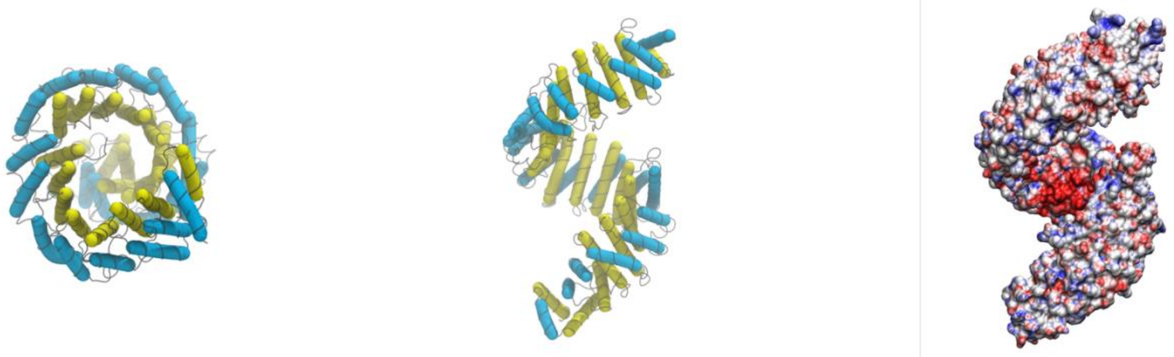

#### TNPO3*

| HEAT1 | HEAT2 | HEAT3 | HEAT4 | HEAT5 | HEAT6 | HEAT7 | HEAT8 | HEAT9 | HEAT10 |
|---|---|---|---|---|---|---|---|---|---|
| 8-20,24-39 | 42-53,57-73 | 81-96,102-118 | 125-134,139-154 | 163-189,195-211 | 223-233,239-255 | 262-285,289-312 | 321-332,343-355 | 359-380,395-410 | 416-426,434-447 |
| **HEAT11** | **HEAT12** | **HEAT13** | **HEAT14** | **HEAT15** | **HEAT16** | **HEAT17** | **HEAT18** | **HEAT19** | **HEAT20** |
| 457-468,475-494 | 499-511,516-530 | 537-546,555-570 | 574-595,608-621 | 633-652,656-672 | 680-694,698-712 | 718-737,746-761 | 772-784,789-803 | 814-844,850-863 | 865-877,892-904 |

*There is an additional A-helix at the end of the sequence [24] (residues 908-920).

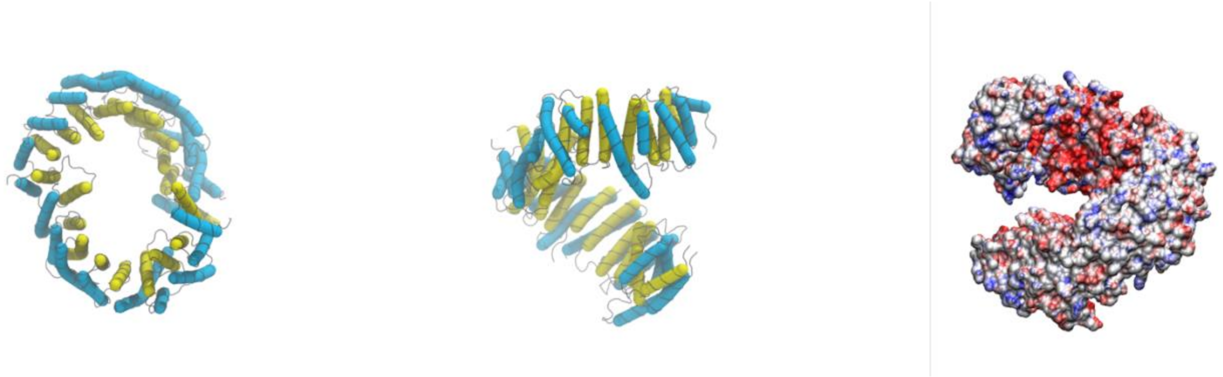

#### Imp5

| HEAT1 | HEAT2 | HEAT3 | HEAT4 | HEAT5 | HEAT6 | HEAT7 | HEAT8 | HEAT9 | HEAT10 |
|---|---|---|---|---|---|---|---|---|---|
| 21-37,41-53 | 56-68,74-90 | 101-117,121-137 | 148-159,163-175 | 187-200,205-221 | 234-248,252-265 | 273-285,291-307 | 316-331,353-368 | 370-386,390-407 | 414-426,431-448 |
| **HEAT11** | **HEAT12** | **HEAT13** | **HEAT14** | **HEAT15** | **HEAT16** | **HEAT17** | **HEAT18** | **HEAT19** | **HEAT20** |
| 450-470,474-490 | 499-521,524-541 | 552-562,569-586 | 594-605,618-633 | 641-652,692-710 | 718-725,735-755 | 757-776,781-798 | 805-833,840-871 | 882-890,896-913 | 924-931,937-954 |
| **HEAT21** | **HEAT22** | **HEAT23** | **HEAT24** | | | | | | |
| 959-974,983-1000 | 1007-1017,1024-1039 | 1052-1062,1074-1087 | 1090-1099,1102-1115 | | | | | | |

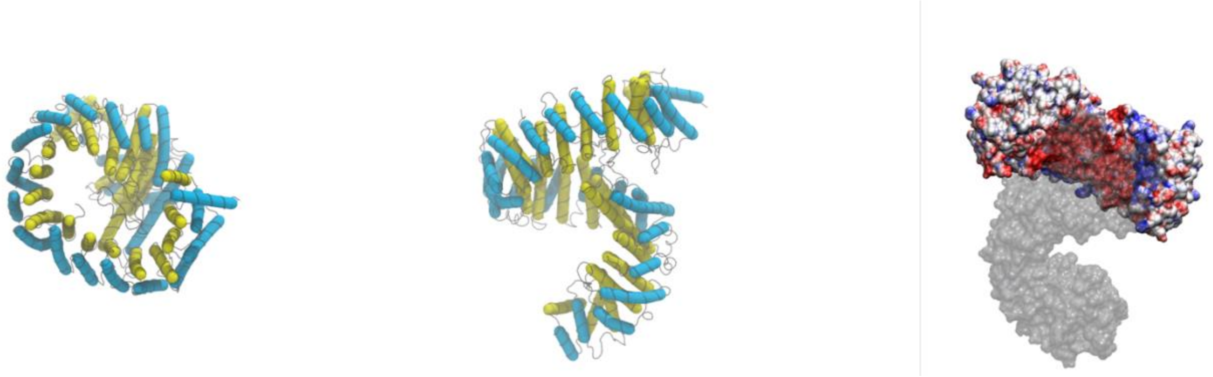

### CRM1

| HEAT1 | HEAT2 | HEAT3 | HEAT4 | HEAT5 | HEAT6 | HEAT7 | HEAT8 | HEAT9 | HEAT10 |
|---|---|---|---|---|---|---|---|---|---|
| - | - | 96-115,125-145 | 148-158,161-180 | 188-215,219-233 | 245-254,260-273 | 280-297,313-339 | 344-358,363-383 | 404-423,449-467 | 469-485,491-503 |
| **HEAT11** | **HEAT12** | **HEAT13** | **HEAT14** | **HEAT15** | **HEAT16** | **HEAT17** | **HEAT18** | **HEAT19** | **HEAT20** |
| 510-530,534-550 | 559-573,580-595 | 610-622,627-642 | 647-674,682-702 | 713-735,743-765 | 769-790,798-811 | 819-835,842-858 | 868-882,887-905 | 908-931,939-954 | 970-985,991-1004 |
| **HEAT21** | | | | | | | | | |
| 1008-1022,1038-1054 | | | | | | | | | |

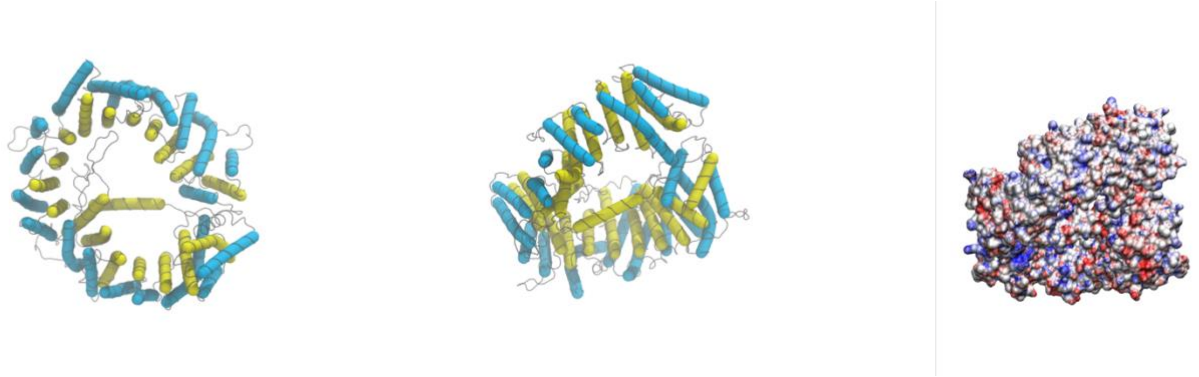

### XPO5*

| HEAT1 | HEAT2 | HEAT3 | HEAT4 | HEAT5 | HEAT6 | HEAT7 | HEAT8 | HEAT9 | HEAT10 |
|---|---|---|---|---|---|---|---|---|---|
| 5-20,27-43 | 46-55,61-77 | 84-100,110-131 | 135-145,147-166 | 172-207,214-236 | 249-257,263-276 | 293-308,313-336 | 348-361,365-380 | 388-405,429-452 | 455-471,498-520 |
| **HEAT11** | **HEAT12** | **HEAT13** | **HEAT14** | **HEAT15** | **HEAT16** | **HEAT17** | **HEAT18** | **HEAT19** | **HEAT20** |
| 528-540,546-565 | 570-582,595-614 | 622-634,642-659 | 662-681,711-734 | 765-787,832-857 | 868-874,885-902 | 911-936,952-976 | 1022-1034,1041-1052 | 1068-1083,1087-1105 | 1111-1116,1123-1134,1146-1150 |

*In the last HEAT repeat, there are two A-helices [25] (residues 1111-1116 and residues 1123-1134). In our contact site analysis and the visualization presented here, the non-helix region between these two helices is considered to be part of the A-helix. Therefore, the last A-helix contains residues 1111-1134.

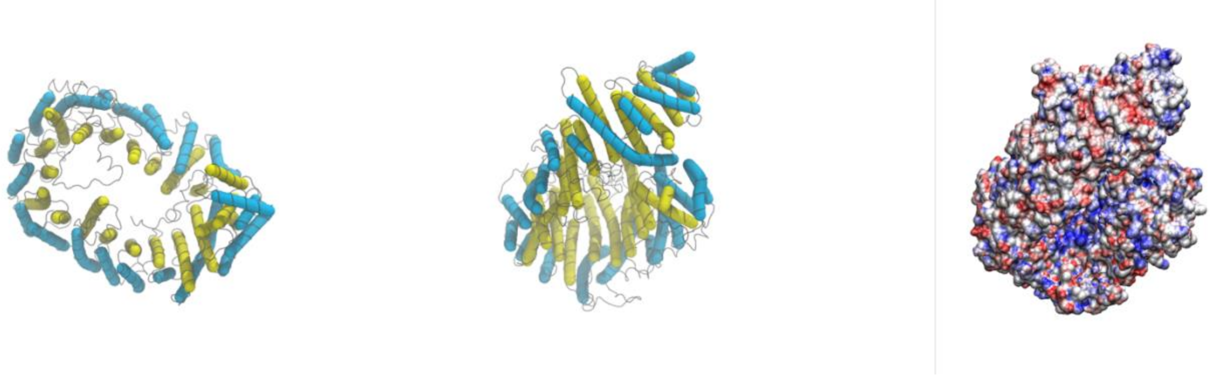

### KAP95

| HEAT1 | HEAT2 | HEAT3 | HEAT4 | HEAT5 | HEAT6 | HEAT7 | HEAT8 | HEAT9 | HEAT10 |
|---|---|---|---|---|---|---|---|---|---|
| 3-15,19-35 | 37-49,55-67 | 90-105,109-126 | 133-143,149-165 | 177-188,195-208 | 219-234,238-255 | 259-275,280-306 | 321-333,347-362 | 367-378,383-395 | 402-418,425-441 |
| **HEAT11** | **HEAT12** | **HEAT13** | **HEAT14** | **HEAT15** | **HEAT16** | **HEAT17** | **HEAT18** | **HEAT19** | |
| 451-463,467-485 | 496-508,516-530 | 535-554,563-586 | 594-607,614-629 | 637-649,654-670 | 678-690,697-713 | 721-733,741-765 | 775-788,796-812 | 825-836,842-860 | |

*In the linker between HEATs 2 and 3, there is a relatively long α-helix with 13 residues.

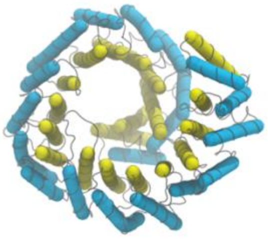
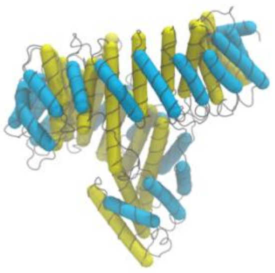
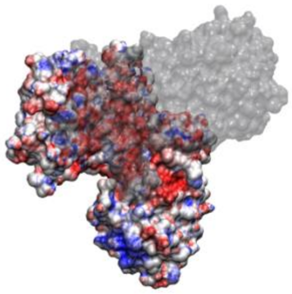

### KAP121

| HEAT1 | HEAT2 | HEAT3 | HEAT4 | HEAT5 | HEAT6 | HEAT7 | HEAT8 | HEAT9 | HEAT10 |
|---|---|---|---|---|---|---|---|---|---|
| 6-19,24-38 | 43-57,61-78 | 96-111,116-128 | 138-149,153-165 | 175-186,191-207 | 220-233,236-253 | 260-273,279-294 | 304-318,343-359 | 364-376,381-395 | 405-413,422-439 |
| **HEAT11** | **HEAT12** | **HEAT13** | **HEAT14** | **HEAT15** | **HEAT16** | **HEAT17** | **HEAT18** | **HEAT19** | **HEAT20** |
| 443-459,465-480 | 490-503,507-524 | 532-545,551-568 | 570-589,596-613 | 621-632,669-689 | 697-709,715-736 | 741-760,764-781 | 788-810,829-849 | 853-869,873-888 | 901-909,914-931 |
| **HEAT21** | **HEAT22** | **HEAT23** | **HEAT24** | | | | | | |
| 935-950,961-978 | 986-994,1002-1018 | 1028-1042,1047-1063 | 1066-1072,1078-1086 | | | | | | |

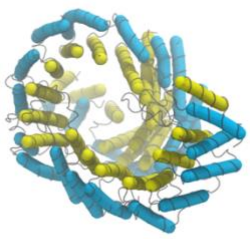
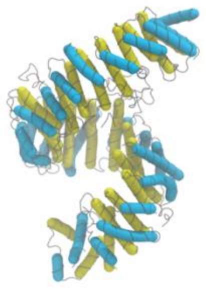
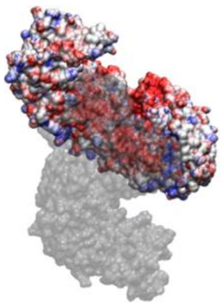

#### KAP114

| **HEAT1** | **HEAT2** | **HEAT3** | **HEAT4** | **HEAT5** | **HEAT6** | **HEAT7** | **HEAT8** | **HEAT9** | **HEAT10** |
|---|---|---|---|---|---|---|---|---|---|
| 2-11,15-31 | 33-45,51-69 | 84-99,105-123 | 129-141,144-157 | 168-181,187-204 | 216-234,243-263 | 271-290,303-321 | 328-342,372-381 | 385-401,408-421 | 431-447,453-470 |
| **HEAT11** | **HEAT12** | **HEAT13** | **HEAT14** | **HEAT15** | **HEAT16** | **HEAT17** | **HEAT18** | **HEAT19** | **HEAT20** |
| 476-494,498-514 | 523-542,546-562 | 571-587,592-606 | 611-635,641-655 | 665-681,685-700 | 710-722,734-743 | 752-766,770-786 | 805-818,823-838 | 879-895,965-979 | 984-990,993-1003 |

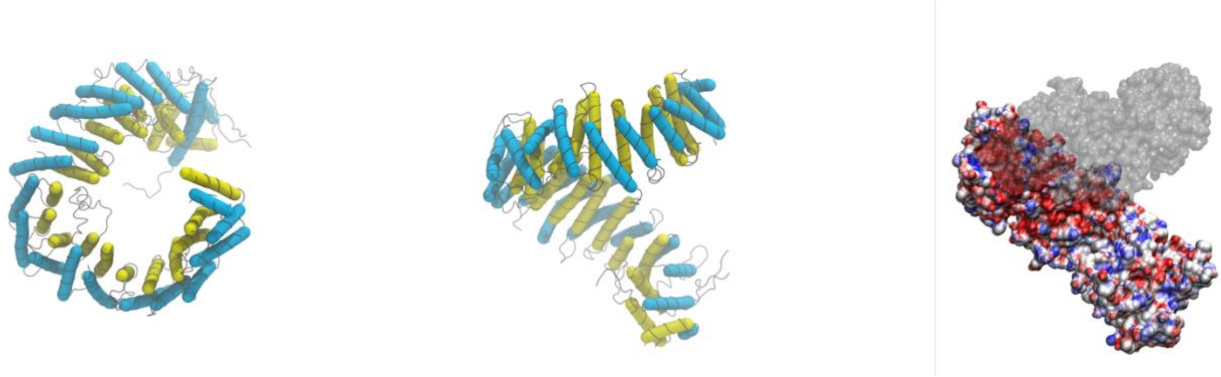

#### KAP124

| **HEAT1** | **HEAT2** | **HEAT3** | **HEAT4** | **HEAT5** | **HEAT6** | **HEAT7** | **HEAT8** | **HEAT9** | **HEAT10** |
|---|---|---|---|---|---|---|---|---|---|
| - | 51-56,64-76 | 84-103,112-129 | 137-147,149-167 | 177-203,207-221 | 233-244,248-260 | 271-290,306-332 | 336-351,356-375 | 421-433,459-478 | 480-496,502-515 |
| **HEAT11** | **HEAT12** | **HEAT13** | **HEAT14** | **HEAT15** | **HEAT16** | **HEAT17** | **HEAT18** | **HEAT19** | **HEAT20** |
| 521-541,545-561 | 570-584,589-611 | 621-633,638-654 | 658-685,693-713 | 717-745,754-776 | 780-801,809-822 | 827-846,853-870 | 879-894,898-918 | 922-944,952-968 | 988-1002,1008-1019 |
| **HEAT21** | | | | | | | | | |
| 1025-1040,1052-1071 | | | | | | | | | |

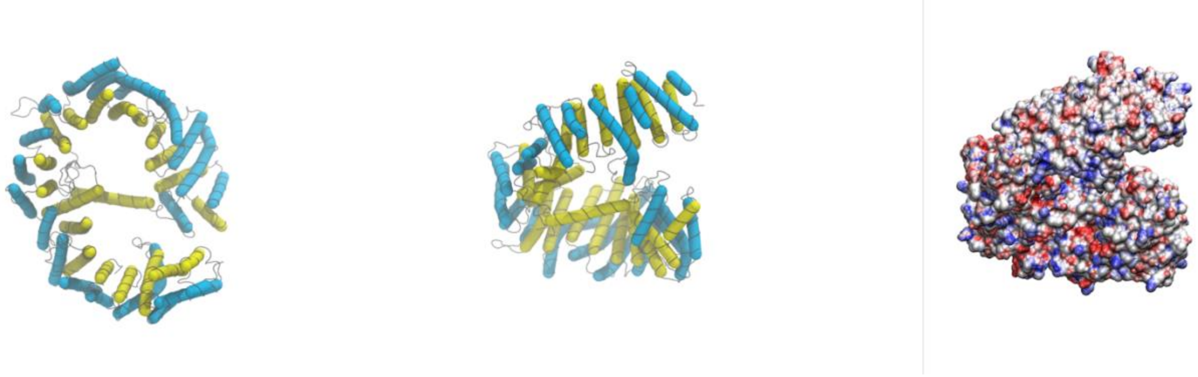

**Table S5** Sequences of Kapβs used for the CG modeling. See also column six of Table S3 for more information about Kapβ sequences.

| NTR name | Amino acid sequence |
|---|---|
| Impβ1 (KPNB1) | MELITILEKTVSPDRLELEAAQKFLERAAVENLPTFLVELSRVLANPGNS<br>QVARVAAGLQIKNSLTSKDPDIKAQYQQRWLAIDANARREVKNYVLHTLG<br>TETYRPSSASQCVAGIACAEIPVNQWPELIPQLVANVTNPNSTEHMKEST<br>LEAIGYICQDIDPEQLQDKSNEILTAIIQGMRKEEPSNNVKLAATNALLN<br>SLEFTKANFDKESERHFIMQVVCEATQCPDTRVRVAALQNLVKIMSLYYQ<br>YMETYMGPALFAITIEAMKSDIDEVALQGIEFWSNVCDEEMDLAIEASEA<br>AEQGRPPEHTSKFYAKGALQYLVPILTQTLTKQDENDDDDDWNPCKAAGV<br>CLMLLATCCEDDIVPHVLPFIKEHIKNPDWRYRDAAVMAFGCILEGPEPS<br>QLKPLVIQAMPTLIELMKDPSVVVRDTAAWTVGRICELLPEAAINDVYLA<br>PLLQCLIEGLSAEPRVASNVCWAFSSLAEAAYEAADVADDQEEPATYCLS<br>SSFELIVQKLLETTDRPDGHQNNLRSSAYESLMEIVKNSAKDCYPAVQKT<br>TLVIMERLQQVLQMESHIQSTSDRIQFNDLQSLLCATLQNVLRKVQHQDA<br>LQISDVVMASLLRMFQSTAGSGGVQEDALMAVSTLVEVLGGEFLKYMEAF<br>KPFLGIGLKNYAEYQVCLAAVGLVGDLCRALQSNIIPFCDEVMQLLLENL<br>GNENVHRSVKPQILSVFGDIALAIGGEFKKYLEVVLNTLQQASQAQVDKS<br>DYDMVDYLNELRESCLEAYTGIVQGLKGDQENVHPDVMLVQPRVEFILSF<br>IDHIAGDEDHTDGVVACAAGLIGDLCTAFGKDVLKLVEARPMIHELLTEG<br>RRSKTNKAKTLARWATKELRKLKNQA |
| TNPO1 | MEYEWKPDEQGLQQILQLLKESQSPDTTIQRTVQQKLEQLNQYPDFNNYL<br>IFVLTKLKSEDEPTRSLSGLILKNNVKAHFQNFPNGVTDFIKSECLNNIG<br>DSSPLIRATVGILITTIASKGELQNWPDLLPKLCSLLDSEDYNTCEGAFG<br>ALQKICEDSAEILDSDVLDRPLNIMIPKFLQFFKHSSPKIRSHAVACVNQ<br>FIISRTQALMLHIDSFIENLFALAGDEEPEVRKNVCRALVMLLEVRMDRL<br>LPHMHNIVEYMLQRTQDQDENVALEACEFWLTLAEQPICKDVLVRHLPKL<br>IPVLVNGMKYSDIDIILLKGDVEEDETIPDSEQDIRPRFHRSRTVAQQHD<br>EDGIEEEDDDDDEIDDDDTISDWNLRKCSAAALDVLANVYRDELLPHILP<br>LLKELLFHHEWVVKESGILVLGAIAEGCMQGMIPYLPELIPHLIQCLSDK<br>KALVRSITCWTLSRYAHWVVSQPPDTYLKPLMTELLKRILDSNKRVQEAA<br>CSAFATLEEEACTELVPYLAYILDTLVFAFSKYQHKNLLILYDAIGTLAD<br>SVGHHLNKPEYIQMLMPPLIQKWNMLKDEDKDLFPLLECLSSVATALQSG<br>FLPYCEPVYQRCVNLVQKTLAQAMLNNAQPDQYEAPDKDFMIVALDLLSG<br>LAEGLGGNIEQLVARSNILTLMYQCMQDKMPEVRQSSFALLGDLTKACFQ<br>HVKPCIADFMPILGTNLNPEFISVCNNATWAIGEISIQMGIEMQPYIPMV<br>LHQLVEIINRPNTPKTLLENTAITIGRLGYVCPQEVAPMLQQFIRPWCTS<br>LRNIRDNEEKDSAFRGICTMISVNPSGVIQDFIFFCDAVASWINPKDDLR<br>DMFCKILHGFKNQVGDENWRRFSDQFPLPLKERLAAFYGV |
| TNPO3 | MEGAKPTLQLVYQAVQALYHDPDPSGKERASFWLGELQRSVHAWEISDQL<br>LQIRQDVESCYFAAQTMKMKIQTSFYELPTDSHASLRDSLLTHIQNLKDL<br>SPVIVTQLALAIADLALQMPSWKGCVQTLVEKYSNDVTSLPFLLEILTVL<br>PEEVHSRSLRIGANRRTEIIEDLAFYSSTVVSLLMTCVEKAGTDEKMLMK<br>VFRCLGSWFNLGVLDSNFMANNKLLALLFEVLQQDKTSSNLHEAASDCVC<br>SALYAIENVETNLPLAMQLFQGVLTLETAYHMAVAREDLDKVLNYCRIFT<br>ELCETFLEKIVCTPGQGLGDLRTLELLLICAGHPQYEVVEISFNFWYRLG |

| | |
|---|---|
| | EHLYKTNDEVIHGIFKAYIQRLLHALARHCQLEPDHEGVPEETDDFGEFR<br>MRVSDLVKDLIFLIGSMECFAQLYSTLKEGNPPWEVTEAVLFIMAAIAKS<br>VDPENNPTLVEVLEGVVRLPETVHTAVRYTSIELVGEMSEVVDRNPQFLD<br>PVLGYLMKGLAEKPLASAAAKAIHNICSVCRDHMAQHFNGLLEIARSLDS<br>FLLSPEAAVGLLKGTALVLARLPLDKITECLSELCSVQVMALKKLLSQEP<br>SNGISSDPTVFLDRLAVIFRHTNPIVENGQTHPCQKVIQEIWPVLSETLN<br>KHRADNRIVERCCRCLRFAVRCVGKGSAALLQPLVTQMVNVYHVHQHSCF<br>LYLGSILVDEYGMEEGCRQGLLDMLQALCIPTFQLLEQQNGLQNHPDTVD<br>DLFRLATRFIQRSPVTLLRSQVVIPILQWAIASTTLDHRDANCSVMRFLR<br>DLIHTGVANDHEEDFELRKELIGQVMNQLGQQLVSQLLHTCCFCLPPYTL<br>PDVAEVLWEIMQVDRPTFCRWLENSLKGLPKETTVGAVTVTHKQLTDFHK<br>QVTSAEECKQVCWALRDFTRLFR |
| Imp5 | AMAAAEQQQFYLLLGNLLSPDNVVRKQAEETYENIPGQSKITFLLQAIRN<br>TTAAEEARQMAAVLLRRLLSSAFDEVYPALPSDVQTAIKSELLMIIQMET<br>QSSMRKKVCDIAAELARNLIDEDGNNQWPEGLKFLFDSVSSQNVGLREAA<br>LHIFWNFPGIFGNQQQHYLDVIKRMLVQCMQDQEHPSIRTLSARATAAFI<br>LANEHNVALFKHFADLLPGFLQAVNDSCYQNDDSVLKSLVEIADTVPKYL<br>RPHLEATLQLSLKLCGDTSLNNMQRQLALEVIVTLSETAAAMLRKHTNIV<br>AQTIPQMLAMMVDLEEDEDWANADELEDDDFDSNAVAGESALDRMACGLG<br>GKLVLPMIKEHIMQMLQNPDWKYRHAGLMALSAIGEGCHQQMEGILNEIV<br>NFVLLFLQDPHPRVRYAACNAVGQMATDFAPGFQKKFHEKVIAALLQTME<br>DQGNQRVQAHAAAALINFTEDCPKSLLIPYLDNLVKHLHSIMVLKLQELI<br>QKGTKLVLEQVVTSIASVADTAEEKFVPYYDLFMPSLKHIVENAVQKELR<br>LLRGKTIECISLIGLAVGKEKFMQDASDVMQLLLKTQTDFNDMEDDDPQI<br>SYMISAWARMCKILGKEFQQYLPVVMGPLMKTASIKPEVALLDTQDMENM<br>SDDDGWEFVNLGDQQSFGIKTAGLEEKSTACQMLVCYAKELKEGFVEYTE<br>QVVKLMVPLLKFYFHDGVRVAAAESMPLLLECARVRGPEYLTQMWHFMCD<br>ALIKAIGTEPDSDVLSEIMHSFAKCIEVMGDGCLNNEHFEELGGILKAKL<br>EEHFKNQELRQVKRQDEDYDEQVEESLQDEDDNDVYILTKVSDILHSIFS<br>SYKEKVLPWFEQLLPLIVNLICPHRPWPDRQWGLCIFDDVIEHCSPASFK<br>YAEYFLRPMLQYVCDNSPEVRQAAAYGLGVMAQYGGDNYRPFCTEALPLL<br>VRVIQSADSKTKENVNATENCISAVGKIMKFKPDCVNVEEVLPHWLSWLP<br>LHEDKEEAVQTFNYLCDLIESNHPIVLGPNNTNLPKIFSIIAEGEMHEAI<br>KHEDPCAKRLANVVRQVQTSGGLWTECIAQLSPEQQAAIQELLNSA |
| XPO1/CRM1 | NMNTKYYGLQILENVIKTRWKILPRNQCEGIKKYVVGLIIKTSSDPTCVE<br>KEKVYIGKLNMILVQILKQEWPKHWPTFISDIVGASRTSESLCQNNMVIL<br>KLLSEEVFDFSSGQITQVKSKHLKDSMCNEFSQIFQLCQFVMENSQNAPL<br>VHATLETLLRFLNWIPLGYIFETKLISTLIYKFLNVPMFRNVSLKCLTEI<br>AGVSVSQYEEQFVTLFTLTMMQLKQMLPLNTNIRLAYSNGKDDEQNFIQN<br>LSLFLCTFLKEHDQLIEKRLNLRETLMEALHYMLLVSEVEETEIFKICLE<br>YWNHLAAELYRESPFSTSASPLLSGSQHFDVPPRRQLYLPMLFKVRLLMV<br>SRMAKPEEVLVVENDQGEVVREFMKDTDSINLYKNMRETLVYLTHLDYVD<br>TERIMTEKLHNQVNGTEWSWKNLNTLCWAIGSISGAMHEEDEKRFLVTVI<br>KDLLGLCEQKRGKDNKAIIASNIMYIVGQYPRFLRAHWKFLKTVVNKLFE<br>FMHETHDGVQDMACDTFIKIAQKCRRHFVQVQVGEVMPFIDEILNNINTI<br>ICDLQPQQVHTFYEAVGYMIGAQTDQTVQEHLIEKYMLLPNQVWDSIIQQ<br>ATKNVDILKDPETVKQLGSILKTNVRACKAVGHPFVIQLGRIYLDMLNVY<br>KCLSENISAAIQANGEMVTKQPLIRSMRTVKRETLKLISGWVSRSNDPQM<br>VAENFVPPLLDAVLIDYQRNVPAAREPEVLSTMAIIVNKLGGHITAEIPQ<br>IFDAVFECTLNMINKDFEEYPEHRTNFFLLLQAVNSHCFPAFLAIPPTQF |

| | |
|---|---|
| | KLVLDSIIWAFKHTMRNVADTGLQILFTLLQNVAQEEAAAQSFYQTYFCD<br>ILQHIFSVVTDTSHTAGLTMHASILAYMFNLVEEGKISTSLNPGNPVNNQ<br>IFLQEYVANLLKSAFPHLQDAQVKLFVTGLFSLNQDIPAFKEHLRDFLVQ<br>IKEFAGEDTSDLFLEEREIALRQADEEKHKRQMSVPGIFNPHEIPEEMCD |
| XPO5 | MAMDQVNALCEQLVKAVTVMMDPNSTQRYRLEALKFCEEFKEKCPICVPC<br>GLRLAEKTQVAIVRHFGLQILEHVVKFRWNGMSRLEKVYLKNSVMELIAN<br>GTLNILEEENHIKDALSRIVVEMIKREWPQHWPDMLIELDTLSKQGETQT<br>ELVMFILLRLAEDVVTFQTLPPQRRRDIQQTLTQNMERIFSFLLNTLQEN<br>VNKYQQVKTDTSQESKAQANCRVGVAALNTLAGYIDWVSMSHITAENCKL<br>LEILCLLLNEQELQLGAAECLLIAVSRKGKLEDRKPLMVLFGDVAMHYIL<br>SAAQTADGGGLVEKHYVFLKRLCQVLCALGNQLCALLGADSDVETPSNFG<br>KYLESFLAFTTHPSQFLRSSTQMTWGALFRHEILSRDPLLLAIIPKYLRA<br>SMTNLVKMGFPSKTDSPSCEYSRFDFDSDEDFNAFFNSSRAQQGEVMRLA<br>CRLDPKTSFQMAGEWLKYQLSTFLDAGSVNSCSAVGTGEGSLCSVFSPSF<br>VQWEAMTLFLESVITQMFRTLNREEIPVNDGIELLQMVLNFDTKDPLILS<br>CVLTNVSALFPFVTYRPEFLPQVFSKLFSSVTFETVEESKAPRTRAVRNV<br>RRHACSSIIKMCRDYPQLVLPNFDMLYNHVKQLLSNELLLTQMEKCALME<br>ALVLISNQFKNYERQKVFLEELMAPVASIWLSQDMHRVLSDVDAFIAYVG<br>TDQKSCDPGLEDPCGLNRARMSFCVYSILGVVKRTCWPTDLEEAKAGGFV<br>VGYTSSGNPIFRNPCTEQILKLLDNLLALIRTHNTLYAPEMLAKMAEPFT<br>KALDMLDAEKSAILGLPQPLLELNDSPVFKTVLERMQRFFSTLYENCFHI<br>LGKAGPSMQQDFYTVEDLATQLLSSAFVNLNNIPDYRLRPMLRVFVKPLV<br>LFCPPEHYEALVSPILGPLFTYLHMRLSQKWQVINQRSLLCGEDEAADEN<br>PESQEMLEEQLVRMLTREVMDLITVCCVSKKGADHSSAPPADGDDEEMMA<br>TEVTPSAMAELTDLGKCLMKHEDVCTALLITAFNSLAWKDTLSCQRTTSQ<br>LCWPLLKQVLSGTLLADAVTWLFTSVLKGLQMHGQHDGCMASLVHLAFQI<br>YEALRPRYLEIRAVMEQIPEIQKDSLDQFDCKLLNPSLQKVADKRRKDQF<br>KRLIAGCIGKPLGEQFRKEVHIKNLPSLFKKTKPMLETEVLDNDGGGLAT<br>IFEP |
| KAP95 | MSTAEFAQLLENSILSPDQNIRLTSETQLKKLSNDNFLQFAGLSSQVLID<br>ENTKLEGRILAALTLKNELVSKDSVKTQQFAQRWITQVSPEAKNQIKTNA<br>LTALVSIEPRIANAAAQLIAAIADIELPHGAWPELMKIMVDNTGAEQPEN<br>VKRASLLALGYMCESADPQSQALVSSSNNILIAIVQGAQSTETSKAVRLA<br>ALNALADSLIFIKNNMEREGERNYLMQVVCEATQAEDIEVQAAAFGCLCK<br>IMSLYYTFMKPYMEQALYALTIATMKSPNDKVASMTVEFWSTICEEEIDI<br>AYELAQFPQSPLQSYNFALSSIKDVVPNLLNLLTRQNEDPEDDDWNVSMS<br>AGACLQLFAQNCGNHILEPVLEFVEQNITADNWRNREAAVMAFGSIMDGP<br>DKVQRTYYVHQALPSILNLMNDQSLQVKETTAWCIGRIADSVAESIDPQQ<br>HLPGVVQACLIGLQDHPKVATNCSWTIINLVEQLAEATPSPIYNFYPALV<br>DGLIGAANRIDNEFNARASAFSALTTMVEYATDTVAETSASISTFVMDKL<br>GQTMSVDENQLTLEDAQSLQELQSNILTVLAAVIRKSPSSVEPVADMLMG<br>LFFRLLEKKDSAFIEDDVFYAISALAASLGKGFEKYLETFSPYLLKALNQ<br>VDSPVSITAVGFIADISNSLEEDFRRYSDAMMNVLAQMISNPNARRELKP<br>AVLSVFGDIASNIGADFIPYLNDIMALCVAAQNTKPENGTLEALDYQIKV<br>LEAVLDAYVGIVAGLHDKPEALFPYVGTIFQFIAQVAEDPQLYSEDATSR<br>AAVGLIGDIAAMFPDGSIKQFYGQDWVIDYIKRTRSGQLFSQATKDTARW<br>AREQQKRQLSL |
| KAP121 | MSALPEEVNRTLLQIVQAFASPDNQIRSVAEKALSEEWITENNIEYLLTF<br>LAEQAAFSQDTTVAALSAVLFRKLALKAPITHIRKEVLAQIRSSLLKGFL |

| | |
|---|---|
| | SERADSIRHKLSDAIAECVQDDLPAWPELLQALIESLKSGNPNFRESSFR<br>ILTTVPYLITAVDINSILPIFQSGFTDASDNVKIAAVTAFVGYFKQLPKS<br>EWSKLGILLPSLLNSLPRFLDDGKDDALASVFESLIELVELAPKLFKDMF<br>DQIIQFTDMVIKNKDLEPPARTTALELLTVFSENAPQMCKSNQNYGQTLV<br>MVTLIMMTEVSIDDDDAAEWIESDDTDDEEEVTYDHARQALDRVALKLGG<br>EYLAAPLFQYLQQMITSTEWRERFAAMMALSSAAEGCADVLIGEIPKILD<br>MVIPLINDPHPRVQYGCCNVLGQISTDFSPFIQRTAHDRILPALISKLTS<br>ECTSRVQTHAAAALVNFSEFASKDILEPYLDSLLTNLLVLLQSNKLYVQE<br>QALTTIAFIAEAAKNKFIKYYDTLMPLLLNVLKVNNKDNSVLKGKCMECA<br>TLIGFAVGKEKFHEHSQELISILVALQNSDIDEDDALRSYLEQSWSRICR<br>ILGDDFVPLLPIVIPPLLITAKATQDVGLIEEEEAANFQQYPDWDVVQVQ<br>GKHIAIHTSVLDDKVSAMELLQSYATLLRGQFAVYVKEVMEEIALPSLDF<br>YLHDGVRAAGATLIPILLSCLLAATGTQNEELVLLWHKASSKLIGGLMSE<br>PMPEITQVYHNSLVNGIKVMGDNCLSEDQLAAFTKGVSANLTDTYERMQD<br>RHGDGDEYNENIDEEEDFTDEDLLDEINKSIAAVLKTTNGHYLKNLENIW<br>PMINTFLLDNEPILVIFALVVIGDLIQYGGEQTASMKNAFIPKVTECLIS<br>PDARIRQAASYIIGVCAQYAPSTYADVCIPTLDTLVQIVDFPGSKLEENR<br>SSTENASAAIAKILYAYNSNIPNVDTYTANWFKTLPTITDKEAASFNYQF<br>LSQLIENNSPIVCAQSNISAVVDSVIQALNERSLTEREGQTVISSVKKLL<br>GFLPSSDAMAIFNRYPADIMEKVHKWFA |
| KAP114 | GPLGSMDINELIIGAQSADKHTREVAETQLLQWCDSDASQVFKALANVAL<br>QHEASLESRQFALLSLRKLITMYWSPGFESYRSTSNVEIDVKDFIREVLL<br>KLCLNDNENTKIKNGASYCIVQISAVDFPDQWPQLLTVIYDAISHQHSLN<br>AMSLLNEIYDDVVSEEMFFEGGIGLATMEIVFKVLNTETSTLIAKIAALK<br>LLKACLLQMSSHNEYDEASRKSFVSQCLATSLQILGQLLTLNFGNVDVIS<br>QLKFKSIIYENLVFIKNDFSRKHFSSELQKQFKIMAIQDLENVTHINANV<br>ETTESEPLLETVHDCSIYIVEFLTSVCTLQFSVEEMNKIITSLTILCQLS<br>SETREIWTSDFNTFVSKETGLAASYNVRDQANEFFTSLPNPQLSLIFKVV<br>SNDIEHSTCNYSTLESLLYLLQCILLNDDEITGENIDQSLQILIKTLENI<br>LVSQEIPELILARAILTIPRVLDKFIDALPDIKPLTSAFLAKSLNLALKS<br>DKELIKSATLIAFTYYCYFAELDSVLGPEVCSETQEKVIRIINQVSSDAE<br>EDTNGALMEVLSQVISYNPKEPHSRKEILQAEFHLVFTISSEDPANVQVV<br>VQSQECLEKLLDNINMDNYKNYIELCLPSFINVLDSNNANNYRYSPLLSL<br>VLEFITVFLKKKPNDGFLPDEINQYLFEPLAKVLAFSTEDETLQLATEAF<br>SYLIFNTDTRAMEPRLMDIMKVLERLLSLEVSDSAAMNVGPLVVAIFTRF<br>SKEIQPLIGRILEAVVVRLIKTQNISTEQNLLSVLCFLTCNDPKQTVDFL<br>SSFQIDNTDALTLVMRKWIEAFEVIRGEKRIKENIVALSNLFFLNDKRLQ<br>KVVVNGNLIPYEGDLIITRSMAKKMPDRYVQVPLYTKIIKLFVSELSFQS<br>KQPNPEQLITSDIKQEVVNANKDDDNDDWEDVDDVLDYDKLKEYIDDDVD<br>EEADDDSDDITGLMDVKESVVQLLVRFFKEVASKDVSGFHCIYETLSDSE<br>RKVLSEALL |
| KAP120 | ASSLNELNLVQVLEQASNPQHIRSDVQKLAEQQLRQWETQAGFHYLLQSI<br>YLNLSNSLQIRWLAVIQFKNGVDKYWRSTRINAIPKDEKASIRGRLFEMI<br>DEQNNQLCIQNAQASARIARLDFPVEWPTLFEDLENLLNDEIIRKDSVKI<br>YNILMHINQIVKVLGTARIGRCRPAMQSKVPLILPLIVRIYLQSFEEWTT<br>SSNLNYEDLSSLQVSYLALKVLRRIICEGYDRPQTDQSVCDFIKLSVSHF<br>EMLISNHENFKKFDIYEKFIKCLGKLYFNLVTGSPANFILLPCSTQILIT<br>YTRLIFDKAPKVYRENSDVTGDFWEQTAIRGLLILKRVINFIHKKGAITL<br>KARSDKLTIDASINKINTEFLNENLITRLVDTLMEWYLRLRPTELENWFM<br>DPEEWINEQMATSYEYQIRPCAENVFQDLMNTFSELLVPYLLKKIENDAS |

| | |
|---|---|
| | KLSNSLDDFLRKDAIYASFQLSASAVSEMVDFDRLLIQVFLPEATNTNIS<br>GDELRIIRRRVALIINEWSTVKCSEESKSLCYKLFTNFLTDEDDKVVLLT<br>TVQTVRTMVDDWNFNKDTFQPFLTENVHLLLRKILPSVSLTETRLYVLNT<br>LSDIIIQTKPLISRDLLVEILQIIPNLWEIATNNASEAILANALLRLLRN<br>LVSSLGSQSHLTWDIAIPVVALACDPSSMQYQLLSEDGYELWGMLLQNFS<br>SHDQEFDDKFVELVPFLKYGIETHTEILPTLLEIIKSYALILNPVDFFSN<br>NTFQDIFKQMSKYLLKLREDSFQLVLEIWEILILSNESDYENLLLQKFYE<br>TGVLSALFDAIFLEEAPSSYLCSQIIQIIARISYVNPDALMTFLATYHDN<br>LPTSNENARMPESIRKIVSKDQTYDSVVNKLLTGWIVCFRDIFDPKFKKV<br>HILGISSLLRTGLVPILTEFSSIASLWIEMLEEINETNRGDCEKYHLNDI<br>VTEQSIAFHPLTAEQLRYHQLCKNNDPVHNISLKDFISQSMEYLESHLGV<br>ERYQEFLKTINPSLLENLQMFLSIQPQEARP |
| KAP124 | AWQKADQILQFSTNPQSKFIALSILDKLITRKWKLLPNDHRIGIRNFVVG<br>MIISMCQDDEVFKTQKNLINKSDLTLVQILKQEWPQNWPEFIPELIGSSS<br>SSVNVCENNMIVLKLLSEEVFDFSAEQMTQAKALHLKNSMSKEFEQIFKL<br>CFQVLEQGSSSSLIVATLESLLRYLHWIPYRYIYETNILELLSTKFMTSP<br>DTRAITLKCLTEVSNLKIPQDNDLIKRQTVLFFQNTLQQIATSVMPVTAD<br>LKATYANANGNDQSFLQDLAMFLTTYLARNRALLESDESLRELLLNAHQY<br>LIQLSKIEERELFKTTLDYWHNLVADLFYEPLKKHIYEEICSQLRLVIIE<br>NMVRPEEVLVVENDEGEIVREFVKESDTIQLYKSEREVLVYLTHLNVIDT<br>EEIMISKLARQIDGSEWSWHNINTLSWAIGSISGTMSEDTEKRFVVTVIK<br>DLLDLTVKKRGKDNKAVVASDIMYVVGQYPRFLKAHWNFLRTVILKLFEF<br>MHETHEGVQDMACDTFIKIVQKCKYHFVIQQPRESEPFIQTIIRDIQKTT<br>ADLQPQQVHTFYKACGIIISEERSVAERNRLLSDLMQLPNMAWDTIVEQS<br>TANPTLLLDSETVKIIANIIKTNVAVCTSMGADFYPQLGHIYYNMLQLYR<br>AVSSMISAQVAAEGLIATKTPKVRGLRTIKKEILKLVETYISKARNLDDV<br>VKVLVEPLLNAVLEDYMNNVPDARDAEVLNCMTTVVEKVGHMIPQGVILI<br>LQSVFECTLDMINKDFTEYPEHRVEFYKLLKVINEKSFAAFLELPPAAFK<br>LFVDAICWAFKHNNRDVEVNGLQIALDLVKNIERMGNVPFANEFHKNYFF<br>IFVSETFFVLTDSDHKSGFSKQALLLMKLISLVYDNKTSNQVYLSQYLAN<br>MLSNAFPHLTSEQIASFLSALTKQYKDLVVFKGTLRDFLVQIKEVGGDPT<br>DYLFAEDKENALMEQNRLEREKAAKIGGLLKPSELDD |

### 7 Supporting references

1. Ghavami, A., E. Van der Giessen, and P.R. Onck, *Coarse-Grained Potentials for Local Interactions in Unfolded Proteins.* Journal of Chemical Theory and Computation, 2013. **9**(1): p. 432-440.

2. Ghavami, A., L.M. Veenhoff, E. Van der Giessen, and P.R. Onck, *Probing the disordered domain of the nuclear pore complex through coarse-grained molecular dynamics simulations.* Biophysical Journal, 2014. **107**(6): p. 1393-1402.

3. Fragasso, A., et al., *A designer FG-Nup that reconstitutes the selective transport barrier of the nuclear pore complex.* Nat Commun, 2021. **12**(1): p. 2010.

4. Jafarinia, H., E. Van der Giessen, and P.R. Onck, *Phase Separation of Toxic Dipeptide Repeat Proteins Related to C9orf72 ALS/FTD.* Biophys J, 2020. **119**(4): p. 843-851.

5. Hutten, S., et al., *Nuclear Import Receptors Directly Bind to Arginine-Rich Dipeptide Repeat Proteins and Suppress Their Pathological Interactions.* Cell Rep, 2020. **33**(12): p. 108538.

6. Crowley, P.B. and A. Golovin, *Cation–π interactions in protein–protein interfaces.* Proteins: Structure, Function, and Bioinformatics, 2005. **59**(2): p. 231-239.

7. Gallivan, J.P. and D.A. Dougherty, *Cation-π interactions in structural biology.* Proceedings of the National Academy of Sciences, 1999. **96**(17): p. 9459-9464.

8. Wang, J., et al., *A Molecular Grammar Governing the Driving Forces for Phase Separation of Prion-like RNA Binding Proteins.* Cell, 2018. **174**(3): p. 688-699.e16.

9. Brady, J.P., et al., *Structural and hydrodynamic properties of an intrinsically disordered region of a germ cell-specific protein on phase separation.* Proceedings of the National Academy of Sciences, 2017. **114**(39): p. E8194-E8203.

10. Krainer, G., et al., *Reentrant liquid condensate phase of proteins is stabilized by hydrophobic and non-ionic interactions.* Nat Commun, 2021. **12**(1): p. 1085.

11. Song, J., S.C. Ng, P. Tompa, K.A. Lee, and H.S. Chan, *Polycation-π interactions are a driving force for molecular recognition by an intrinsically disordered oncoprotein family.* PLoS Comput Biol, 2013. **9**(9): p. e1003239.

12. Miyazawa, S. and R.L. Jernigan, *Estimation of effective interresidue contact energies from protein crystal structures: quasi-chemical approximation.* Macromolecules, 1985. **18**(3): p. 534-552.

13. Kim, D.E., D. Chivian, and D. Baker, *Protein structure prediction and analysis using the Robetta server.* Nucleic Acids Res, 2004. **32**(Web Server issue): p. W526-31.

14. Saito, N. and Y. Matsuura, *A 2.1-Å-resolution crystal structure of unliganded CRM1 reveals the mechanism of autoinhibition.* J Mol Biol, 2013. **425**(2): p. 350-64.

15. McGuffin, L.J., K. Bryson, and D.T. Jones, *The PSIPRED protein structure prediction server.* Bioinformatics, 2000. **16**(4): p. 404-5.

16. Higurashi, M., T. Ishida, and K. Kinoshita, *PiSite: a database of protein interaction sites using multiple binding states in the PDB.* Nucleic Acids Res, 2009. **37**(Database issue): p. D360-4.

17. Frishman, D. and P. Argos, *Knowledge-based protein secondary structure assignment.* Proteins, 1995. **23**(4): p. 566-79.

18. Cingolani, G., J. Bednenko, M.T. Gillespie, and L. Gerace, *Molecular basis for the recognition of a nonclassical nuclear localization signal by importin beta.* Mol Cell, 2002. **10**(6): p. 1345-53.

19. Imasaki, T., et al., *Structural basis for substrate recognition and dissociation by human transportin 1.* Mol Cell, 2007. **28**(1): p. 57-67.

20. Zhang, Z.C. and Y.M. Chook, *Structural and energetic basis of ALS-causing mutations in the atypical proline-tyrosine nuclear localization signal of the Fused in Sarcoma protein (FUS).* Proc Natl Acad Sci U S A, 2012. **109**(30): p. 12017-21.

21. Soniat, M. and Y.M. Chook, *Karyopherin-β2 Recognition of a PY-NLS Variant that Lacks the Proline-Tyrosine Motif.* Structure, 2016. **24**(10): p. 1802-1809.

22. Jang, S., et al., *Differential role for phosphorylation in alternative polyadenylation function versus nuclear import of SR-like protein CPSF6.* Nucleic Acids Res, 2019. **47**(9): p. 4663-4683.

23. Cingolani, G., C. Petosa, K. Weis, and C.W. Müller, *Structure of importin-β bound to the IBB domain of importin-α.* Nature, 1999. **399**(6733): p. 221-229.

24. Maertens, G.N., et al., *Structural basis for nuclear import of splicing factors by human Transportin 3.* Proc Natl Acad Sci U S A, 2014. **111**(7): p. 2728-33.

25. Yamazawa, R., et al., *Structural Basis for Selective Binding of Export Cargoes by Exportin-5.* Structure, 2018. **26**(10): p. 1393-1398.e2.
